# Tumor Agnostic Drug Delivery with Self-Agglomerating Nanohydrogels (SANGs)

**DOI:** 10.1101/2024.01.22.575714

**Authors:** Stephen N. Housley, Sebinne Lee, Lilya V. Matyunina, Olivia A. Herrmann, Minati Satpathy, Johana C. Arboleda, John F. McDonald, M.G. Finn

## Abstract

RNA interference (RNAi) holds unique potential as a clinically viable modality to pharmacologically regulate oncogenes in sequence-specific manner. Despite its potential, systemic delivery of RNAi to tumors encounters myriad obstructions and strategies to overcome barriers have largely consisted of academic demonstrations, with few approaches reaching patients. Here, we report the development of a self-agglomerating nanohydrogel (SANGs) platform that is efficiently internalized by cancer cells, is agnostic to RNAi payload, and achieves functional suppression of multiple oncogene targets. After intravenous injection, SANGs preferentially accumulated and were retained ubiquitously in primary and metastatic loci in three aggressive cancer models in a species-agnostic manner. SANGs efficiently delivered multiple RNAi payloads that significantly suppressed oncogene expression and sensitized previously resistance tumors *in vivo*. SANGs were found to be safe and well tolerated in simulated clinical applications across three species. We then propose and verify a novel emergent mechanism by which SANGs achieve durable solid-tumor delivery without direct functionalization. Overall, our SANGs platform is an enabling technology for RNAi-based cancer therapeutics and is poised for advanced pharmaceutical development with multiple solid-tumor indications.

**One-Sentence Summary:** Our nanostructure achieves safe and durable tumor-agnostic delivery through a newly described environmentally-responsive mechanism.

## Main

RNA interference (RNAi) uses small interfering RNA (siRNA) and microRNA (miRNA) to silence expression of target genes in a sequence-specific manner(*1, 2*). RNAi molecules hold unique potential as clinically viable modalities for manifold diseases(*3, 4*). Oncological indications are of particular interest because the vast majority cancer-promoting genes are “undruggable”(*4*). In order to reach its potential, RNAi molecules must be delivered to tumors, preferably after systemic injection. Such delivery is challenging due to rapid clearance, nuclease susceptibility, and inability to traverse plasma membranes, resulting in insufficient delivery to and inadequate penetration of tumors(*3, 5, 6*).

Numerous strategies have been developed to overcome these barriers, dominated by complexation with cationic polymers or inclusion within liposomes to form lipid nanoparticles (LNPs)(*7-11*). While these strategies have improved delivery, trafficking is poorly understood and LNPs have drawbacks(*12-15*). Notably, the vast majority of an LNP dose becomes trapped in the liver and is not efficiently taken up by cancerous cells after systemic injection(*16-18*). This leaves the majority of cancers, especially metastasis, untreated(*16*). Cationic polymers face different challenges: they bind nonspecifically to many cells types(*19*) and interact with serum components, resulting in short circulation times and toxic side effects. Surface shielding by PEGylation improves stability, reduces nonspecific binding and clearance, and prolongs circulation, it also impairs target cell uptake(*20, 21*). While ligand functionalization can improve specificity, it increases manufacturing complexity and costs, reduces stability, and requires prior knowledge of the cell type of interest(*16, 17*). Collectively, the development of a systemic, tumor agnostic delivery system that is effective, safe, and simple to manufacture remains an important missing link for therapeutic translation of RNAi.

Here we report the development of a self-agglomerating nanohydrogel (SANGs) platform that achieves durable solid-tumor delivery without direct functionalization. We demonstrate that SANGs enable potent delivery of RNAi molecules *in vitro* and exhibit preferential accumulation and retention in primary and metastatic loci in three cancer models in a species-agnostic manner. SANGs efficiently delivered multiple RNAi payloads, sensitizing previously drug-resistant tumors. SANGs were found to be safe and well tolerated in simulated clinical applications across three species. We also provide a proposed mechanism by which SANGs operate, which takes advantage of an emergent property of these materials *in vivo*.

### SANGs synthesis and characterization

We previously observed that polyacrylamide-based nanohydrogels functionalized with tumor-targeting peptides exclusively targeted ovarian tumors(*22*). These core-shell nanohydrogels are made from inexpensive materials and are easily formulated as soft and biocompatible particles using a simple, two-stage free-radical precipitation polymerization method (Fig. 1a and Supplementary Fig. 1)(*23-25*). However, instead of functionalizing the shell layer with directing ligands, we explored the capabilities of untargeted nanohydrogels (SANGs), chemically functionalized only by the attachment of a small amount of fluorophore by amide bond formation. Physicochemical properties were characterized (Fig. 1b-c and Supplementary Figs. 2, 3) and independently validated by the National Characterization Laboratory under a variety of solvent, pH, dilutions, and temperature conditions (Fig. 1d and Supplementary Fig. 4). Both siRNA-loaded and unloaded SANGs were found to be unchanged in size, morphology, and concentration upon storage at -25°C, 4°C, and 25°C out to 60 days in aqueous conditions (Fig. 1e). SANGs prepared with negative control (NC) and scrambled RNAi molecules were shown to have no significant effect on the viability of HEY-A8-F8 cells (75 nM in particles, Fig. 1f).

**Figure 1.**
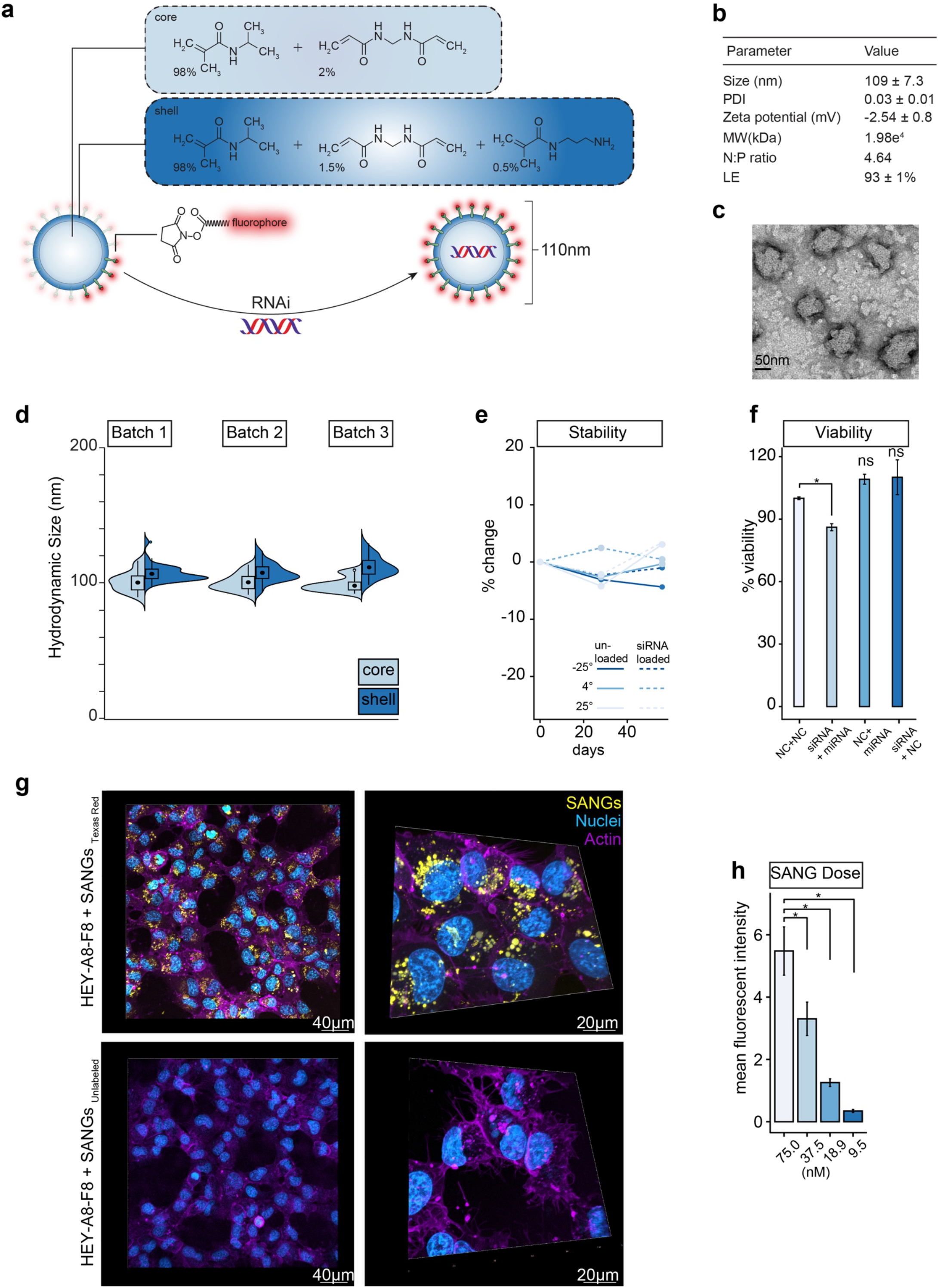
Physicochemical property characterization of SANG and its cellular uptake. (**a**) A schematic diagram of the protocol for the synthesis and preparation of fluorescently labeled siRNA-loaded SANGs. (**b**) Physicochemical property characterization of SANGs, which includes the hydrodynamic size, polydispersion index (PDI), zeta potential, molecular weight (MW in kDa), N:P ratio, and loading efficiency (L.E.). (**c**) Negative-stain TEM image of SANGs. (**d**) reproducibility of SANG production across 3 independent batches showing the complete size distribution of the core and shell components prior to final cleaning and purification.(**e**) Chemical stability of siRNA loaded and unloaded SANGs at -25°C, 4°C and 25°C. (**f**) Cell viabilities (Hey-A8-F8) of SANGs (75nM) differentially loaded with negative control (NC) siRNA-EGFR (siRNA) and mir-429 (miRNA), normalized to saline treated cells. (**g**) Cellular uptake in Hey-A8-F8 ovarian cancer cells following overnight incubation (18 h) with Texas-Red labeled (top) and unlabeled (bottom) SANGs (yellow pseudo-colored for contrast) for both low (left, 20x) and high (right, 63x) magnification views. Cells (purple) visualized with Lectin DyLight™ 649 (Vector Laboratories). (**h**) Quantitative analysis of the dose-dependent mean fluorescent intensity (MFI) measured in Hey-A8-F8 ovarian cancer cells 18 h following incubation Texas-Red labeled SANGs. Data are presented as mean ± sd. (*) indicates statistically significant differences between experimental groups as empirically derived from hierarchical Bayesian model.

### SANGs undergo endosomal uptake and escape

SANGs were taken up readily by HEY-A8-F8 in dose- and time-dependent fashion (Fig. 1g-h and Supplementary Fig. 5). Application of SANGs for increasing lengths of time (ranging from 30 minutes to 24 hours) revealed detectable intracellular uptake by 1 hour, which increased until maximal fluorescent signals were reached at 18 hours (Supplementary Fig. 5). Internalization was comparable for epithelial (OVCAR3) and mesenchymal (HEY-A8-F8) type ovarian cancer cells, and MCF7 breast cancer cells (Supplementary Fig. 6). Internalized SANGs remained unchanged in terms of cellular localization and fluorescence intensity for at least 24 hours (Supplementary Fig. 5-6). SANG cellular uptake was shown to be energy- and clathrin-dependent (Fig.2a-c, Supplementary Fig. 7) and unperturbed by inhibition of caveolar and macropinocytotic mechanisms (Fig. 2c, Supplementary Fig. 7), suggesting a clathrin-mediated endosomal pathway. This conclusion was further corroborated by live-cell imaging with an endosomal marker (wheat germ agglutinin, WGA)(*26, 27*)) and SANGs containing dye-labelled siRNA, showing simultaneous co-localization of all components in HEY-A8-F8 cells at early time points (Supplementary Fig. 8). After 18 hours, however, siRNA and SANGs were observed to be distributed throughout the cytoplasm (Fig. 2d-f and Supplementary Fig. 9), suggesting an efficient process of endosomal escape. At this time point, low correlation (R=0.142±0.16, n=24) between siRNA and endosomes was observed, indicating robust release from the particles (Fig. 2g); at six hours, siRNA and endosomes were much more co-located (Fig. 2h).

**Figure 2.**
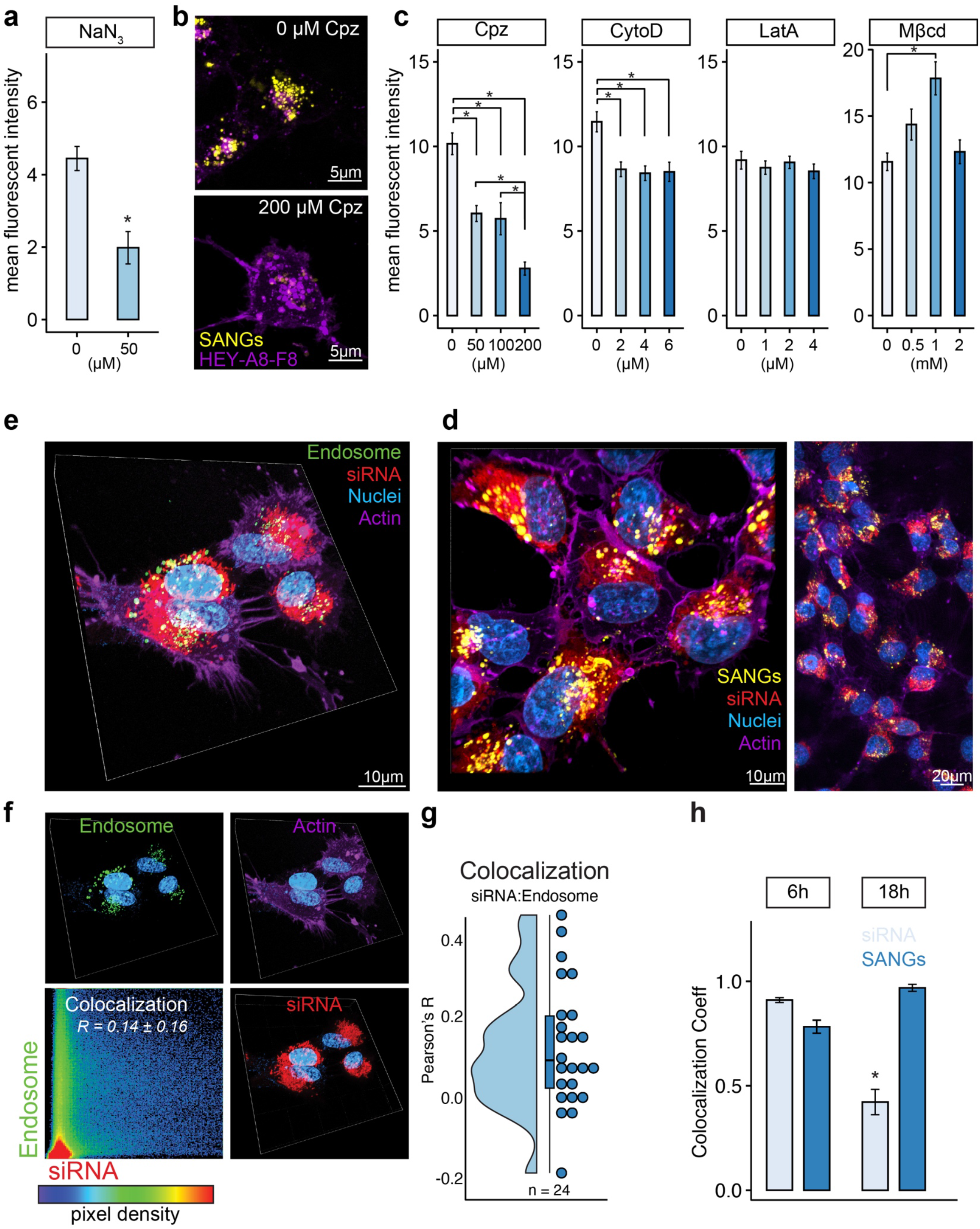
Mechanism of internalization, endosomal escape, and delivery of RNAi payloads. (**a**) Sodium azide (NaN3) dependent reduction of cellular uptake as measured by mean fluorescence intensity. *n = 3* replicates of n = 10 cells. (**b**) Representative confocal fluorescence image (maximum intensity projections from z-stack) showing suppressed SANG internalization by clathrin-mediated endocytosis inhibitor (chlorpromazine [Cpz]). (**c**) Quantification of intracellular mean fluorescent intensity of SANGs following exposure to different doses of macropinocytosis inhibitors (latrunculin A [Lat]; cytochalasin D [CytD]), a caveolae-mediated inhibitor (methyl-β-cyclodextrin [Mβcd]) and Cpz. SANGs were transfected at a final concentration of 75nM. (**d**) Maximum intensity projections from z-stack (10 µm) confocal images of Hey-A8-F8 cells 18 h following incubation with Cy3-siRNA-loaded SANGs (Alexa Fluor 488) at both low (right, 20x) and high (left, 63x) magnifications showing intracellular distribution. (**e**) Three-dimensional rendering of confocal images dual-labeling endosomes (CellLight™ Endosomes-GFP) and siRNA (Cy3) 18 h following Hey-A8-F8 incubation with siRNA-loaded SANGs. (**f**) Individual fluorescent channels shown for clarity. (**f**, lower left) Results of colocalization analysis performed on a pixel-by-pixel basis between endosomes and siRNA channels, where every pixel is plotted based on its intensity level. Scatterplot colors represent pixel density. (**g**) Quantitative results of colocalization analysis summarized by the Pearson’s correlation coefficients (R) between endosomes and siRNA. *n = 24* cellular replicates. (**h**) siRNA release from SANGs quantified by colocalization coefficients (mean ± sd) between SANGs loaded with Cy3-labeled siRNA at 6 and 18 hours after incubation with Hey-A8-F8 cells. *n = 24* cellular replicates. SANGs were transfected at a final particle concentration of 75 nM particle concentration. (*) indicates statistically significant differences between experimental groups as empirically derived from hierarchical Bayesian model (stan_glm): 95% highest density intervals do not overlap between groupwise contrasts.

Consistent with the apparent escape of siRNA to the cytosol, efficient suppression of relevant proteins was observed. Thus, EGFR production by overexpressing HEY-A8-F8 cells was diminished by more than 60% 12 hours after delivery of 20 nM siRNA in SANG particles (Supplementary Fig. 10). The same formulation substituting the appropriate siRNA induced similar knockdown of KRAS, Glut1, and miRNA-429, all known regulators of cancer cell proliferation, migration, and invasion. SANGs containing scrambled siRNA had no effect. The degree of reduction in protein expression was similar to that observed using the same concentration of siRNA delivered with lipofectamine (Supplementary Fig. 10).

### SANG biodistribution and retention in mouse and rat

The *in vivo* biodistribution of SANGs was explored with xenograft and orthotopic murine models of ovarian and breast cancer and a genetically-engineered rat model of colorectal cancer to survey different tumor types, induction methods, species, contribution of the immune system. All cancer strains employed were engineered to express luciferase and thus could be readily followed by bioluminescence imaging (BLI).

Within minutes following i.v. injection in murine ovarian and breast cancer models, SANGs accumulated in regions with higher BLI signal indicating presumptive tumoral targeting (Fig. 3a and Supplementary Fig. 11). *In vivo* distribution of SANGs was distinct from unloaded siRNA and free dyes, which were detected in regions devoid of BLI signal in lungs (within 5 min) and in liver shortly thereafter (30 min) (Fig. 3a). By 24h, the specificity of biodistribution improved out to 72 h (Fig. 3b) where little (if any) SANGs were detected outside BLI inferred tumors. SANGs remained closely associated with BLI signals 7 days after i.v. delivery indicating preferential retention in tumors.

**Figure 3.**
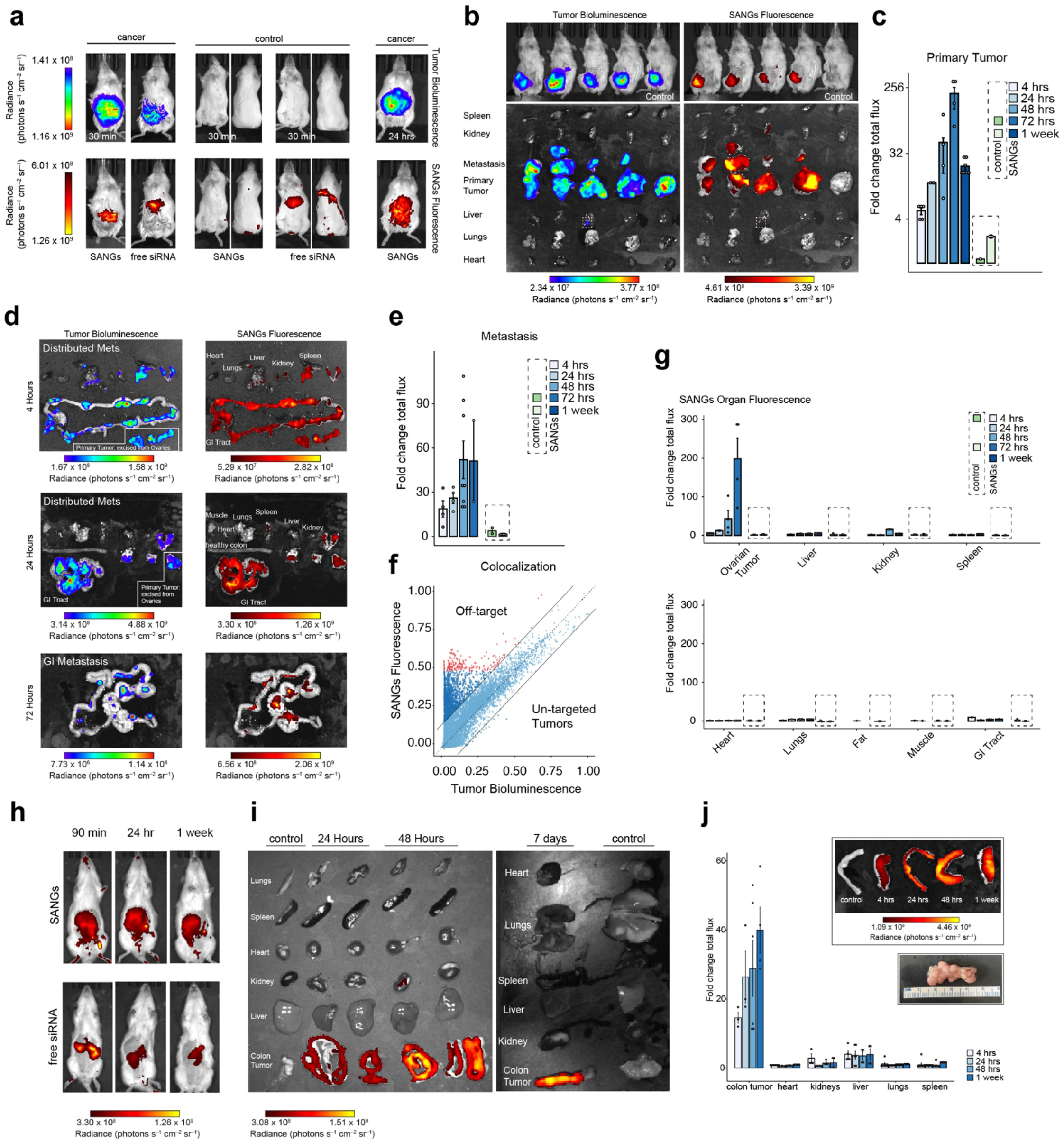
Tumor agnostic, cross species *in vivo* and *ex vivo* biodistribution. (**a**) Whole-body imaging of the mice after intravenous injection of free siRNA or SANGs (1 mg•kg^−1^). Tumor bioluminescence and SANGs fluorescence were imaged sequentially in cancer bearing and control mice showing the immediate (30 min) and early distribution (24 hours). (**b**) Whole-body imaging of cancer-bearing mice 72 hours after intravenous injection of SANGs (1 mg•kg^−1^) or saline control (top row). Ex vivo imaging of tumors (Hey-A8-F8) and major organs immediately imaged after in vivo imaging was completed. (**c**) Quantification of SANG delivery and retention to primary tumors, normalized against SANG fluorescence signal in heart. Intravenous delivery of free siRNA used as quantitative control. (**d**) Representative ex vivo images at indicated time points after intravenous injection of SANGs (1 mg•kg^−1^) to mice following metastatic tumor induction (typically 18-21days). (**e**) Quantification of SANG delivery and retention to metastatic tumors, normalized against SANG fluorescence signal in heart. Intravenous delivery of free siRNA used as quantitative control. (**f**) Colocalization analysis of *ex vivo* SANG fluorescence and tumor bioluminescence. Dotted line of identity indicates perfect colocalization with ±15% bounding conditions (light blue). Upper quadrant shows low (dark blue) and high-intensity (red) off-target SANGs. Lower quadrant shows low (dark blue) and high-intensity (orange) untargeted cancer. (**g**)Quantification of SANG delivery and retention across specified time points across tumors, liver, kidney, spleen, heart, lungs, visceral fat, skeletal muscle and GI tract all normalized against SANG fluorescence signal in heart. Intravenous delivery of free siRNA used as quantitative control. (**h**) Whole-body imaging of Pirc rats after intravenous injection of free siRNA or SANGs (1 mg•kg^−1^). SANGs fluorescence were imaged sequentially in cancer bearing showing the immediate (90 min), early distribution (24 hours), and retention (1 week). (**i**) Ex vivo imaging of tumors and major organs and (**j**) quantification of the specificity of SANG delivery and retention to tumors and major organs (liver, kidney, spleen, heart, lungs), normalized against SANG fluorescence signal in heart. Top inset shows enlarged colorectal tumor biodistribution of SANGs across time. Representative color photograph of a section the internal lumen of the descending colon of a Pirc rat indicating diffuse adenocarcinomas. Data are presented as mean ± sd. (*) indicates statistically significant differences between experimental groups as empirically derived from hierarchical Bayesian model (stan_glm): 95% highest density intervals do not overlap between groupwise contrasts.

Analysis of excised tissues confirmed SANGs rapidly targeted tumors, reaching 5.1±1.2-fold increase by 4 hours and a maximum (198-fold) at 72 hours Fig. 3c. While significantly reduced compared to peak, we find SANGs were retained in primary tumors 7 days after systemic delivery (Fig. 3c). In contrast, intravenous infusion of free siRNA (20 µM) gave significantly lower tumor localization at 3 and 7 days (1.14±0.02 and 2.29±0.17 fold above background) (Fig. 3c)(*28*).

To test the metastatic targeting capacity of SANGs, a limitation of existing RNAi delivery platforms(*29*), we modeled late-stage ovarian cancer by IP administration of HEY-A8-F8 cells. After confirming tumor induction and extensive abdominal metastasis, we i.v. administered SANGs and quantified biodistribution as described above. We observed expansive tumor load across the spleen, GI-tract, liver, and occasionally epidural metastasis (Supplementary Fig. 12). Surprisingly, we also observed rapid (4h) and highly specific accumulation of SANGs in diffuse metastatic lesions. The temporal biodistribution in metastatic loci closely mirrored that of primary tumors independent of organ while signal in the non-cancerous portions of organs remained largely devoid of SANG signal (Fig. 3e and Supplementary Fig. 11-12).

Colocalization analysis of *ex vivo* SANG fluorescence and tumor bioluminescence (Fig. 3e) showed strong association between SANGs and diffuse metastases (R=0.84, *p*<0.001, Supplementary Fig. 13) with >96% of the SANG signal residing within 15% of the line of identity. Few if any (<0.1%) metastatic loci were untargeted by SANGs (below line of identity) while only <3.5% of SANGs were detected in regions without any BLI confirmed metastasis. These results indicate a low level of off-target accumulation and corroborate biodistribution data observed in less metastatic murine models. Taken together, these results indicate that SANGs rapidly target both primary and metastatic murine tumors, are retained for extended periods of time, and have minimal off-target accumulation (Fig. 3g).

Intravenous delivery of SANGs to rats genetically engineered to spontaneously develop colorectal cancer resulted in immediate diffuse SANG signal across the abdominal cavity which began to coalesce in small punctate intensities around the perimeter of the abdomen, suggestive of preferential targeting to the diffuse tumors (Fig. 3h and Supplementary Fig. 14). SANG distribution remained localized in this fashion for 7-days with increasing signal-to-noise ratio over time, indicating preferential tumor retention. In contrast, free siRNA distributed mostly to the liver followed by rapid elimination (Fig. 3h). *Ex vivo* analysis of colon with diffuse adenocarcinomas (Fig. 3j) showed detectable SANG signal at 4 hours, increased signal at 24 and 48 hours, and a maximal recorded value at 7 days (Fig. 3i-j); mouse biodistribution peaked at 72 h (Fig. 3c). As in mouse, SANGs remained largely absent from other major rat organs with the exception of a nominal increase detected in liver and a transient increase (at 4 hours) in the kidney (Fig. 3i-j).

*In vivo* pharmacokinetics and clearance studies revealed an apparent blood distribution half-life of approximately 2.5 h for SANGs and an elimination half-life of ∼13.4 h (Supplementary Fig. 15) in rat. While this value is comparable to that of other nanoformulations(*28, 30*), SANGs behave differently in at least one important respect. We found relatively high SANG concentrations in blood at 24 and 48 h with between 2.8-5.6% of the initial dose remaining in circulation, a factor that likely played a role in the favorable tumor biodistribution and retention. Analysis of urine and feces showed rapid and near complete (∼92%) excretion of naked siRNA by 24 h which was dominated by renal metabolism whereas only 16% of the SANG dose was excreted during the first 24 h. These data show that a large fraction of SANGs leaves systemic circulation rapidly, partitions efficiently to tumors, and perhaps is also sequestered in low concentrations in tissue such as fat, to be released over time.

### In vivo functionality of SANGs

When examined by confocal fluorescence microscopy, SANGs were found distributed throughout ovarian carcinoma (HEY-A8-F8) tumor sections and deep within tumor parenchyma, exhibiting an extravasated distribution into the tumor interstitium (Fig. 4a and Supplementary Fig. 16). A similar result was observed in mice with orthotopically implanted ovarian carcinomas (HEY-A8-F8) and breast cancers (MDA-MB-231) (Supplementary Fig. 17) as well as in rats with advanced colorectal cancer, where we saw strong colocalization with EGFR-overexpressing tumor cells (Fig. 4f). In all cased, the extravasated distribution was retained for at least 7 days.

**Figure 4.**
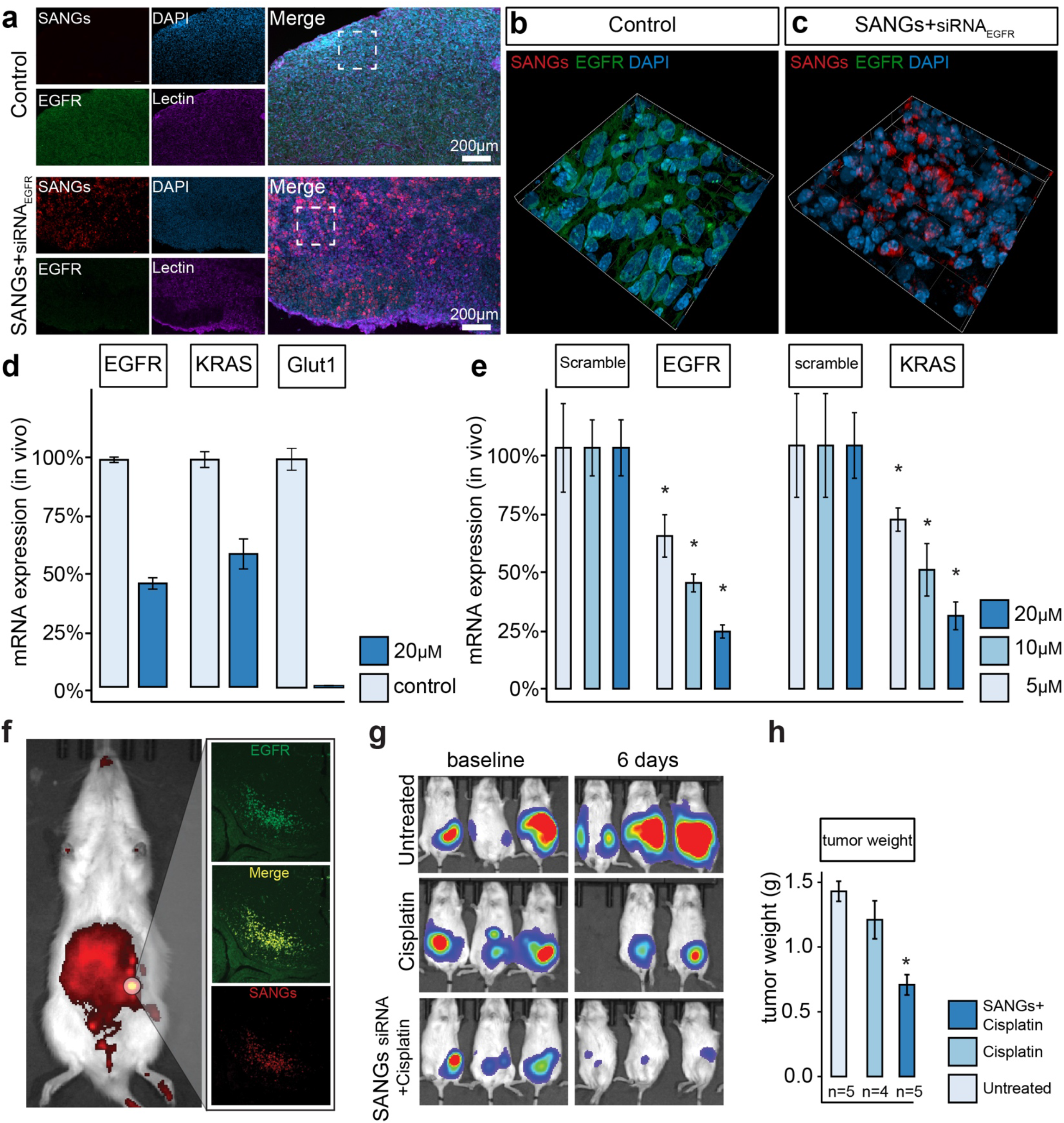
*In vivo* tumor penetration and efficacy. (**a**) representative 20x LSCM images of tumors sections (orthotopic Hey-A8-F8) from mice 48 hours after intravenous administration of either negative control siRNA loaded SANGs (top row) or siRNA against EGFR loaded SANG (bottom row). Three-dimensional (z-stack) confocal microscopic images show expanded view (63x) of white dotted sections (**a**). Intracellular distribution of endosomes and siRNA 18 hours following Hey-A8-F8 incubation with SANGs loaded with Cy3-labeled siRNA. Merged images of cells nuclei (DAPI), EGFR (Alexa Fluor 488) and SANGs (Texas Red). (**d**) Down-regulation of EGFR, KRAS, and Glut1 mRNA levels as quantified by RT-qPCR (n = 3 independent experiments) in Pirc rats. (**e**) Dose-dependent down-regulation of EGFR and KRAS mRNA levels as quantified by RT-qPCR (n = 3 independent experiments) in tumor bearing mice. The relative level of mRNA expression was calculated over the NC siRNA control each of which was normalized to GAPDH (mice) or RPS18 (rats). (**f**) Whole-body *in vivo* imaging of representative Pirc rat after 1 week after intravenous injection of SANGs (1 mg•kg^−1^) with corresponding confocal microscopic images of EGFR overexpressing colorectal cancer cells and SANGs near-infrared fluorescence demonstrating tumor penetration and retention behavior. (**g**) Whole-body *in vivo* bioluminescence imaging OC tumor–bearing mice assessing tumor load prior to and 6 days after cohorts of animals were treated with saline (n=5), cisplatin (n=4), or siRNA against EGFR loaded SANGs (n=5). Quantification of wet tumor weights following last *in vivo* imaging. Data are presented as mean ± sd. (*) indicates statistically significant differences between experimental groups as empirically derived from hierarchical Bayesian model (stan_glm): 95% highest density intervals do not overlap between groupwise contrasts.

Across murine ovarian (HEY-A8-F8) and rat colorectal cancers, we found that single intravenous infusion of SANGs loaded with siRNA against EGFR, KRAS, or GLUT1 resulted in dose-dependent reduction in mRNA expression (Fig. 4d-e) and protein expression (Fig. 4a-c) in tumor tissue. Therapeutic efficacy of SANG-siRNA was then tested in mice with tumors established from HEY-A8-F8 ovarian carcinoma cells. Enhanced expression of EGFR by these cells is correlated with drug resistance, and is a well-established model(*31, 32*). We treated groups of mice with SANGs loaded with mir-429 and siRNA against EGFR 24 hours prior to cisplatin or cisplatin alone. Six days after treatment, dramatic improvements relative to drug alone were evident from significant inhibition of tumor growth (Fig. 4g) and reduction in tumor weight (Fig. 4h). These data indicate that SANGs successfully extravasate, penetrate tumor microenvironments, gain access to the specific cells of interest in a species- and tumor-agnostic manner, and sensitize previously resistant tumors *in vivo*.

### Systemic delivery of SANGs is minimally toxic

A variety of tests revealed no significant toxicity associated with high doses of either empty or scrambled miRNA/siRNA-loaded SANGs. These included observations with outbred mice (Supplementary Fig. 17a,b) and NOD-SCID mice with ovarian tumors including delivery of cisplatin (Supplementary Fig. 18). A repeat-dose tolerability study of high-dose SANG-siRNA and co-infused oxaliplatin in rats with advanced colorectal cancer(*33*) produced no mortality or any clinical signs of distress while weights recovered back to baseline by study completion (Supplementary Fig. 19). A maximal tolerable dose study in rats revealed only minor changes in blood chemistry at 6 and 24 hours (Supplementary Fig. 17c) and no difference from controls in histopathology of major organs for the three escalating doses (Supplementary Fig. 20). Finally, a single-acute dose study in an adult female (73 kg) Yucatan swine showed no clinically meaningful deviances out to 6 hours following i.v. infusion of SANGs-siRNA at the same high dose (7mg•kg^−1^) with the exception of a transient increase in AST at 5 minutes (Supplementary Fig. 17d). Thus, SANGs induce minimal, if any, toxicity in a variety of animals and experimental conditions. Taken together, these data establish a wide preliminary safety profile and strongly support the systemic use of SANG to deliver RNAi to solid tumor cancers.

### Cancer specific delivery achieved through emergent self-agglomerating mechanism

The results described here set SANGs apart from most nanoparticulate delivery vehicles, their most striking property being selective homing to tumors without the use of designed ligands for cell-surface markers. Four complimentary methods were used to probe the mechanism(s) responsible for this i*n vivo* performance. We sought to distinguish between mechanisms involving cell binding and physical properties of the particles that may promote selective distribution to tumor tissue. First, TEM was used to measure the size and number of SANG particles in randomly-selected fixed-size views of 3 µL samples of SANGs at concentrations ranging from 25.5 nM to 1.6 µM prepared in DI water (Fig. 5a). We also modeled the number of SANGs predicted to be in a given micrographic view (Fig. 5b). Results revealed an interesting dynamic behavior. At the lowest three concentrations studied, the number and size distribution of SANGs closely matched the predicted values (Fig. 5b). At higher SANG concentrations, however, this relationship no longer held. Instead, progressively smaller numbers of discriminable SANGs were observed, with increasing size distributions (Fig. 5b and Supplementary Fig. 21). High magnification views showed different representative forms of tightly clustered SANGs that emerge above 156 nM. Similarly, aqueous dynamic light scattering (DLS) on two different instruments showed strongly monodispersed hydrodynamic size distribution at low (52 nM) concentrations, but clear evidence of larger and less regular aggregates at 156 nM (Fig. 5c and Supplementary Fig. 22).

**Figure 5.**
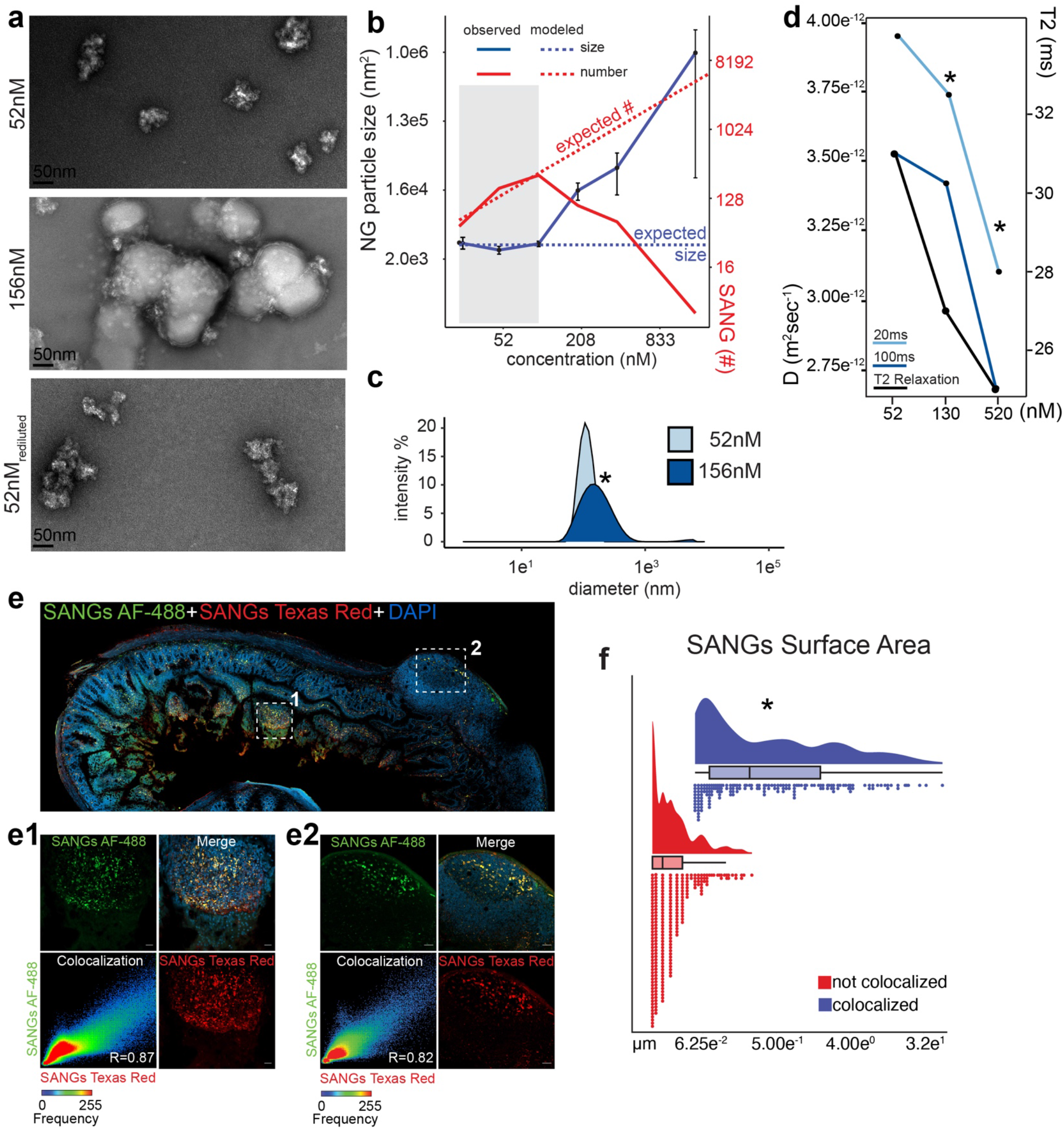
Discovering tumor agnostic mechanism. (**a**) representative phase contrast transmission electron microscopic (TEM) images show SANG behavior at 52nM and 156nM concentrations and following re-dilution challenge taking 156nM sample back to 52nM. (**b**) quantification of complete TEM concentration range studied showing expected (modeled: dotted lines) number and size distribution derived from known concentration and field of view (area). Solid lines and show mean ± sd of observed data across the range of studied concentrations. (**c**) Hydrodynamic size distributions of SANGs at 52nM and 156nM concentrations. (**d**) Diffusion coefficients derived from diffusion-ordered NMR (DOSY) for 20ms and 100ms both pulse sequences and T2 relaxation times across a 10-fold concentration range. (**e**) Representative 20x LSCM tiled image of colon with diffuse adenocarcinomas characterized by marked dysplasia in crypts and villus (box 1) and that penetrate the smooth muscle layers (box 2) showing merged view of both fluorescently labeled SANG populations and DAPI. (**e1-2**) High resolution (63x) maximal projection confocal microscopic images show expanded view of white dotted sections (**e**). Scatter plot of pixel-based colocalization analysis of two representative areas (Costes randomization based colocalization R=0.85±0.02, *p*<2.2^-16^, n=3 animals, 3 sections each). (**f**) Size distribution of colocalized or not-colocalized SANGs derived from object-based analysis of 63x views (n=3 animals, 3 sections). (*) indicates statistically significant differences between experimental groups as empirically derived from hierarchical Bayesian model (stan_glm): 95% highest density intervals do not overlap between groupwise contrasts.

Parallel diffusion-ordered NMR measurements allowed us to detect diffusional changes caused by aggregation. A significant decrease in diffusion coefficient (21.5%) and T2 relaxation time was observed over a 10-fold range of SANG concentration (Fig. 5d and Supplementary Fig. 23). The former parameter indicates slower particle movement in solution and the latter reflects restricted molecular motion(*34, 35*), both consistent with nanogel agglomeration in solution caused at least in part by cross-linking between SANG particles as concentration increases. When concentrated SANGs samples were re-diluted, much of the material returned to the original size range while a portion remained in a larger aggregated state (indicated by DLS and TEM, Fig. 5a and Supplementary Fig. 22).

To test if nanogel aggregation occurs *in vivo*, we intravenously infused two populations of SANGs conjugated to different fluorophores in rats with advanced colorectal cancer. If SANGs do not agglomerate *in vivo*, fluorescent signals from both populations would co-localize at a nominal rate and frequency determined by chance. Colocalization to a significantly greater degree would indicate that agglomeration does indeed occur. Fluorescent signals from SANGS of both colors were readily detected (expanded views, Fig. 5e1-2). Colocalization analysis (Fig. 5e) indicated both SANG populations colocalized significantly greater than chance (Costes randomization-based colocalization R=0.85±0.02, *p*<0.001, n=3 animals, 3 sections each) and were significantly larger deposits (∼74-fold) than SANGs of a single color (Fig. 5f).

## Discussion

Formulated to contain positively charged groups to facilitate the carrying of oligonucleotides, these nanogels were shown to efficiently package and stabilize RNAi molecules and to deliver siRNA and miRNA to three cancer models, equally well in both mouse and rat across primary and metastatic loci. Delivery sensitized drug-resistant tumors, allowing subsequent delivery of chemotherapeutic agents to have a dramatically enhanced effect in arresting tumor growth *in vivo*. The nanogel platform was found to be minimally toxic and well tolerated in mice, rats, and swine in a variety of simulated clinical applications. We are unaware of a delivery platform of any form – polymeric, liposomal, proteinacious, or viral – with this combination of advantages.

The most remarkable properties exhibited by SANGs are: 1) preferential targeting of primary and metastatic tumors(*36*) relative to healthy tissue, 2) prolonged tumor retention, 3) extravasation into the tumor interstitium, gaining access to cancerous cells, and 4) delivery of sufficient RNAi payloads to silence oncogene mRNA and protein expression and result in efficient tumor growth suppression. We believe that the first two of these result from the concentration-dependent “self-agglomeration” behavior demonstrated *in vitro* and *in vivo*. This property allows SANGs to respond to the unique vasculature found in the tumor microenvironment. While altered vasculature in solid tumors has long been suggested to allow drug delivery by the enhanced permeability and retention (EPR) effect, prior results have been muted. We suggest that after SANGs find their way to tumor tissue, its sluggish blood flow (secondary to tortuous nature and elevated resistance(*37, 38*)) allow time for SANG particles to interact, coalesce(*39, 40*) and become entrained in tumors. This responsive behavior enhances the EPR effect and achieves tumor delivery, penetration, and retention to the entire tumor parenchyma regardless of carcinoma type(*6, 41, 42*).

Overall, environmentally-responsive nanostructures of the SANG type may represent an important therapeutic paradigm shift for systemic delivery of RNAi to solid tumors, as it exhibits exceptional *in vivo* performance and substantial therapeutic index that qualify it for advanced development in preparation for clinical applications. The delivery of other molecular payloads may also be envisioned, if the advantageous functional dynamics of SANGs is retained in such cases.

## Acknowledgements

We would like to acknowledge the substantial help of the animal support staff at the two institutions: Georgia Institute of Technology Physiological Research Laboratory (PRL) and Global Center for Medical Innovation T3 Labs. We would like to acknowledge the pathology core facilities at the Yerkes National Primate Research Center for histology and tissue preparation work.

## Funding

McCallum Early Career Fellowship (S.N.H.)

Northside Hospital Foundation, Inc. (S.N.H.)

Ovarian Cancer Institute (J.F.M. and S.N.H.)

Deborah Nash Endowment (J.F.M.)

Georgia Research Alliance (J.F.M. and S.N.H.)

## Author contributions

Conceptualization: S.N.H., M.G.F.

Data curation: S.N.H.

Formal Analysis: S.N.H., L.V.M.

Funding acquisition: S.N.H., J.F.M.

Investigation: S.N.H., L.V.M., S.L., O.A.H., J.C.A., M.S.

Methodology: S.N.H., M.G.F.

Project administration: S.N.H.

Supervision: S.N.H., M.G.F.

Validation: S.N.H., L.V.M., S.L., O.A.H.

Visualization: S.N.H., M.G.F. Writing – original draft: S.N.H.

Writing – review & editing: S.N.H., J.F.M., M.G.F.

## Competing interests

S.N.H and J.F.M have a provisional patent filing (Application No. 63/517,215) related to this work. S.N.H and J.F.M. are cofounders of OnCuRNA. This study could affect their personal financial statuses. The terms of these arrangements have been reviewed and approved by The Georgia Institute of Technology in accordance with its conflict-of-interest policies. All other authors declare no competing interests.

## Data and materials availability

All data, code, and materials used in the analysis are available in the main text or the supplementary materials, in the public repository https://github.com/nickh89/sangs_repo.git and https://figshare.com/account/home#/projects/179187

## Supplementary Materials

Materials and Methods

Figs. S1 to S23

Movies S1 to S3

