## Supplemental Figures Legends for "Tumor Agnostic Drug Delivery with Self-Agglomerating Nanohydrogels (SANGs)"

**Supplementary Figure 1**. **Synthesis and loading.** A schematic diagram of the protocol for the synthesis and preparation of fluorescently labeled siRNA-loaded SANGs. PDI, polydispersity index; DLS, dynamic light scattering.

**Supplementary Figure 2. Physiochemical properties of SANGs. (a)** H nuclear magnetic resonance (NMR) spectra of SANGs in $D_2O$ at 52, 130, and 520nM concentration. Dynamic light scattering distributions of empty (**b**) and siRNA loaded SANGs (**c**). Zeta potential and distributions of SANGs after siRNA loading.

**Supplementary Figure 3. Loading characteristics.** Empirically derived siRNA concentration curve for the quantification of encapsulation and loading efficiency.

**Supplementary Figure 4. National Characterization Laboratory (NCL) Data.** Physiochemical properties, stability, and contamination results (endotoxin) from independent validation at NCL.

**Supplementary Figure 5. Dose- and time-dependent internalization. (top row)** Representative merged confocal fluorescence microscopy images (63x) of Hey-A8-F8 ovarian cancer cells (white) after 18 hours of incubation as function of SANG concentration (red). **(middle row)** Individual SANG fluorescence channel. **(bottom row)** Representative merged confocal fluorescence microscopy images (20x) of Hey-A8-F8 ovarian cancer cells (white) as a function of SANG incubation time (red).

**Supplementary Figure 6. SANG internalization in a variety of cell types.** representative confocal fluorescence microscopy images (63x) of MCF7 breast cancer, OVCAR3 ovarian cancer, and IOSE epithelial ovarian cells following overnight incubation (18 h) with Texas-Red labeled (top) SANGs.

**Supplementary Figure 7. Dose-dependent internalization inhibitors.** Quantification of intracellular mean fluorescent intensity of SANGs following exposure to different doses of clathrin-mediated endocytosis inhibitor (chlorpromazine [Cpz]) macropinocytosis inhibitors (latrunculin A [Lat]; cytochalasin D [CytD]), and a caveolae-mediated inhibitor (methyl-β-cyclodextrin [Mβcd]). Maximum intensity projections from z-stack (63x) confocal images of Hey-A8-F8 cells 18 h following incubation with SANGs. SANGs were transfected at a final concentration of 75nM. Data are presented as mean ± sd. (*) indicates statistically significant differences between experimental groups as empirically derived from hierarchical Bayesian model.

**Supplementary Figure 8. Endosomal uptake and release of siRNA.** Merged super-resolution confocal(63x) confocal images of Hey-A8-F8 cells 18 h following incubation with SANGs loaded with fluorescently labeled siRNA (**right**). Individual fluorescent channels (**left**). Endosomes labeled with wheat germ agglutinin (WGA) a lectin derivative. WGA was applied for 15 minutes prior to imaging. Data reveal a subpopulation of SANGs were actively internalized during WGA incubation (white arrows) while other subpopulations remained devoid of WGA signal (red arrows). * indicate siRNA distributed throughout cytoplasm.

**Supplementary Figure 9. SANGs escape endosomes.** Three-dimensional rendering of confocal images dual-labeling endosomes (CellLight™ Endosomes-GFP) and SANGs (Texas-red) 18 h following Hey-A8-F8 incubation. (**f**) Individual endosome and SANG channels shown for clarity. siRNA release from SANGs quantified by colocalization coefficients (mean ± sd) between SANGs loaded with Cy3-labeled siRNA at 6 and 18 hours after incubation with Hey-A8-F8 cells. $n = 24$ cellular replicates. SANGs were transfected at a final particle concentration of 75 nM particle concentration. Surface renderings of SANGs and Cy3-siRNA channels in two representative cells studied at 6 and 18 hours after incubation. (*) indicates statistically significant differences between experimental groups as empirically derived from hierarchical Bayesian model (stan_glm): 95% highest density intervals do not overlap between groupwise contrasts.

**Supplementary Figure 10. *In vitro* functionality.** (**a**) representative confocal fluorescence microscopy images (20x) show suppression of EGFR protein expression levels in Hey-A8-F8 cells 48 hours after siRNA against EGFR loaded SANG (top row) as compared to negative control siRNA loaded SANGs (bottom row). *n = 3* replicates. In both instances, cells were incubated with SANGs for 12 hours after which media was replaced for remainder of the study. (**b**) Down-regulation of EGFR, KRAS, ZEB1, and Glut1 mRNA levels as quantified by RT-qPCR. The relative level of mRNA expression was calculated over the NC siRNA control. *n = 3* replicates. (**c**) phenotypic shift of canonically mesenchymal Hey-A8-F8 ovarian cancer cells to epithelia phenotype following 12 hours incubation with SANGs loaded with mir-429. Scalebar, 10μM. Data are presented as mean ± sd. (*) indicates statistically significant differences between experimental groups as empirically derived from hierarchical Bayesian model (stan_glm): 95% highest density intervals do not overlap between groupwise contrasts.

**Supplementary Figure 11. Biodistribution comparison across models.** (**a**) Whole-body imaging of the mice 24 h after intravenous injection of SANGs (1 mg•kg$^{-1}$). Tumor bioluminescence and SANGs fluorescence were imaged sequentially in breast cancer bearing mice (orthotopically implanted MDA-MD-231). (**b**) Whole-body imaging of cancer-bearing mice 72 hours after intravenous injection of SANGs (1 mg•kg$^{-1}$). (**c**) Ex vivo imaging of tumors (MDA-MB-231) and major organs immediately imaged after in vivo imaging was completed.

**Supplementary Figure 12. Epidural metastasis spinal cord** (**a**) Ex vivo imaging of the brain and spinal cord from an ovarian cancer (Hey-A8-F8) bearing mouse 72 h after intravenous injection of SANGs (1 mg•kg$^{-1}$). Tumor bioluminescence and SANGs fluorescence were imaged sequentially.

**Supplementary Figure 13. Met targeting quantification.** (**a**) Representative ex vivo images of Tumor bioluminescence and SANGs fluorescence from ovarian cancer (Hey-A8-F8) bearing mouse following metastatic tumor induction (typically 18-21days) 4 h after intravenous injection of SANGs (1 mg•kg$^{-1}$) as shown if Figure 3. (**b**) Image masks utilized for colocalization analysis. (**c**) Computed distributions of colocalization among BLI and SANG channels following 1000 iterations of Costes randomization (blue) and observed colocalization (red line) with one representative shuffled image shown (**d**). Scatter plot of pixel-based colocalization analysis (Costes randomization based colocalization R=0.85±0.02, $p$<2.2$^{-16}$, n=3 animals, 3 sections each).

**Supplementary Figure 14. Time-dependent internalization in rats with advanced colorectal cancer.** Whole-body imaging of Pirc rats after intravenous injection of free siRNA or SANGs (1 mg•kg$^{-1}$). SANGs fluorescence were imaged sequentially at indicated times in Figure 3 in cancer bearing showing

**Supplementary Figure 15. Pharmacokinetics and elimination.** Empirical quantification of SANG concentration in blood (blood) along with bi-exponential model fit (red) after single bolus injections in SD rats. Dotted (purple) and solid (yellow) lines shows siRNA concentration and background fluorescence over time. Inset shows urine and feces elimination of naked siRNA and SANG dose during the first 24 h. Data are presented as mean ± sd (n= 3 rats per group).

**Supplementary Figure 16. Tumor Penetration. (left)** Representative merged confocal fluorescence microscopy image (20x) and expanded (63x of dotted white box) of advanced colorectal tumors, 72 hours after i.v. injection of SANGs (red).

**Supplementary Figure 17. SANGs are minimally toxic.** (**a**) Percentage change of starting weight in response to two i.v. 7mg•kg$^{-1}$ doses of either (1) empty NG or (2) NG loaded with siRNA or miRNA administered to female CD-1 mice (5 animals/group) separated by 24hours. The error bands represent ± 95% CI. (**b**) Complete blood chemistry metrics 14 days after the final dose in CD-1 mice. (**c**) Complete blood chemistry metrics at 6 and 24 hours after i.v. administered of SANGs loaded with siRNA against EGFR at 0, 7, 12, 17 mg•kg$^{-1}$ (3 animals/sex/dose) in Sprague-Dawley rats. Serial blood chemistry metrics (0, 5min, 15min, 30min, 1hr, 2hr,

3hr, 4hr 5hr, and 6hr) after a single bolus i.v. injection of SANGs loaded with siRNA against EGFR ($7mg \cdot kg^{-1}$) in female Domestic Yorkshire Crossbred Swine (6 month, 73kg). Data are presented as mean ± sd unless otherwise noted. (*) indicates statistically significant differences between experimental groups as empirically derived from hierarchical Bayesian model (stan_glm): 95% highest density intervals do not overlap between groupwise contrasts.

**Supplementary Figure 18. SANG toxicity in NOD scid.** Hey-A8-F8 ovarian tumor-bearing mice received the treatments as described in Supplementary Figure 17. Serum levels of ALP, BUN and IL6 at the completion of the therapy study. Mouse weights were measured once every 2 days. Data are presented as mean ± sd (n= 5 mice per group).

**Supplementary Figure 19. SANG toxicity in Pirc rats.** Rats with advanced colorectal cancer (Pirc) received a modeled anticipated clinical treatment schedule. SANG-siEGFR ($7mg \cdot kg^{-1}$) was infused twice weekly for 5 weeks along with a co-infusion of oxaliplatin ($5mg \cdot kg^{-1}$). Weights were measured twice weekly as a surrogate measure for tolerability. Data are presented as mean ± sd (n= 5 mice per group).

**Supplementary Figure 20 Histopathology.** Histology of spleen, kidney, and liver in SD rats 24 hours after randomly assigned to receive saline control, or escalating SANG-siEGFR doses 7, 13, or $17mg \cdot kg^{-1}$ as described in Supplementary Figure 17. Shown are representative images from 4 independent experiments.

**Supplementary Figure 21 TEM. (a)** Fixed view negative-stain TEM images as a function of the SANG concentration and corresponding particle analysis masks (**b**). high magnification SEM of gold coated SANGs.

**Supplementary Figure 22 SANG concentration dependent size change.** Hydrodynamic sizes of SANGs at sequentially escalating sample concentrations followed by re-dilution (grey inset). First (light blue) and second (dark blue) dynamic light scattering peak data are presented as mean ± sd (n=3).

**Supplementary Figure 23 DOSY Raw data.**
Attenuation of the NMR signals during pulsed field gradient experiments. Diffusion coefficients derived from best fit curves. Measured T1-T2 correlations as a function of the SANG concentration, obtained by DOSY with $D_20$ and SANG distributions labeled.
