## Supplemental Figures for "Tumor Agnostic Drug Delivery with Self-Agglomerating Nanohydrogels (SANGs)"

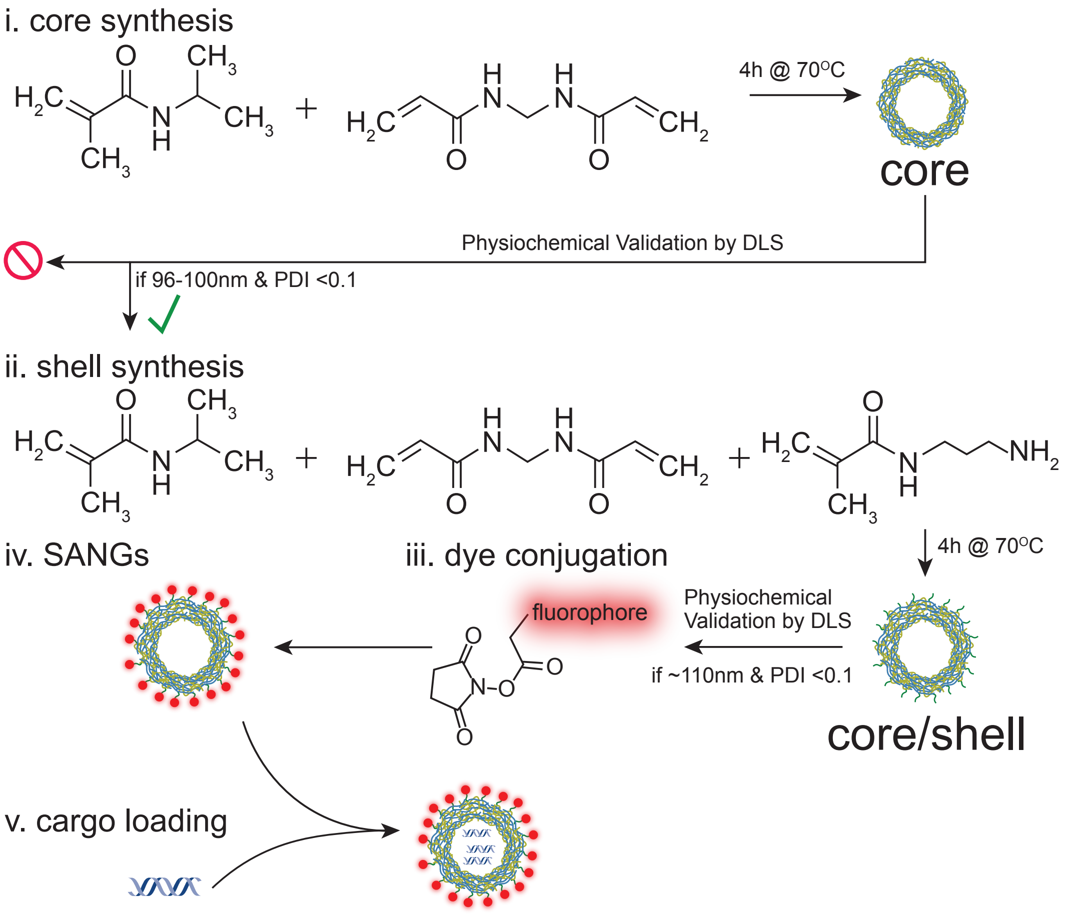

i. core synthesis
4h @ 70°C
core
Physiochemical Validation by DLS
if 96-100nm & PDI <0.1
ii. shell synthesis
4h @ 70°C
iv. SANGs
iii. dye conjugation
fluorophore
Physiochemical
Validation by DLS
if ~110nm & PDI <0.1
core/shell
v. cargo loading

**a**

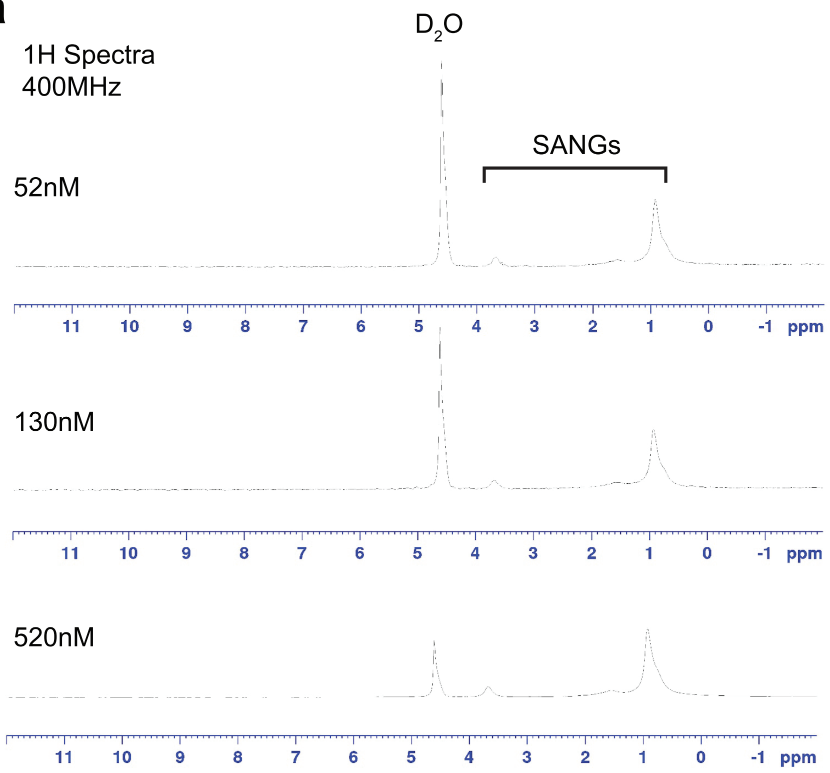


**b**

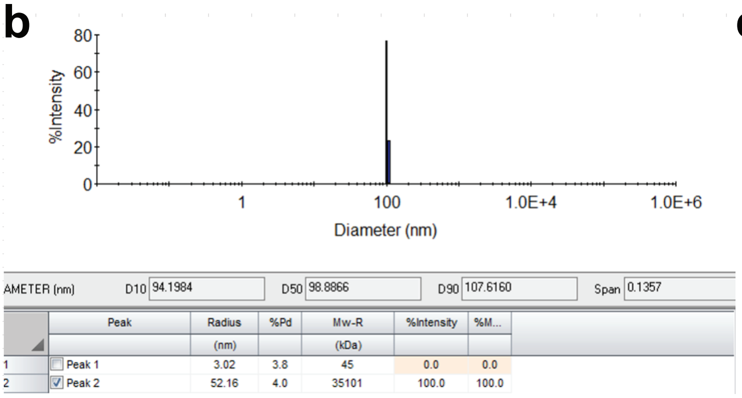


SANGs- empty in 5% saline

**c**

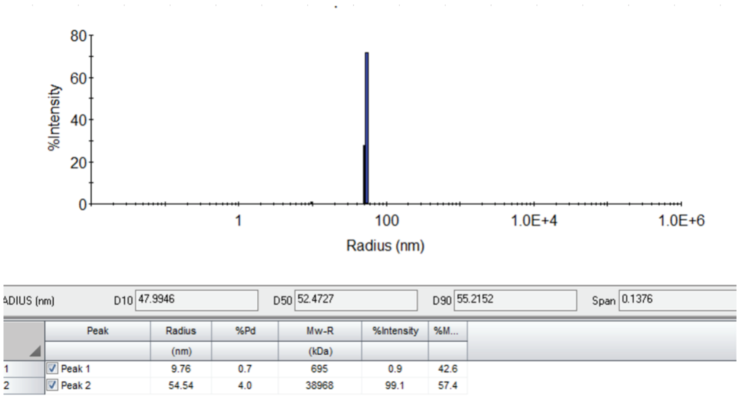


SANGs-siRNA$_{EGFR}$ 5% saline

**d**

| | SANG concentration | Radius z-avg | PDI | Zeta-potential (mv) | SD |
|---|---|---|---|---|---|
| SANGs-siRNA$_{nc}$ | 52nM | 54.27 | 0.0266 | - 1.61 | 0.36 |
| SANGs-siRNA$_{glut1}$ | 52nM | 54.47 | 0.020 | - 1.89 | 0.82 |

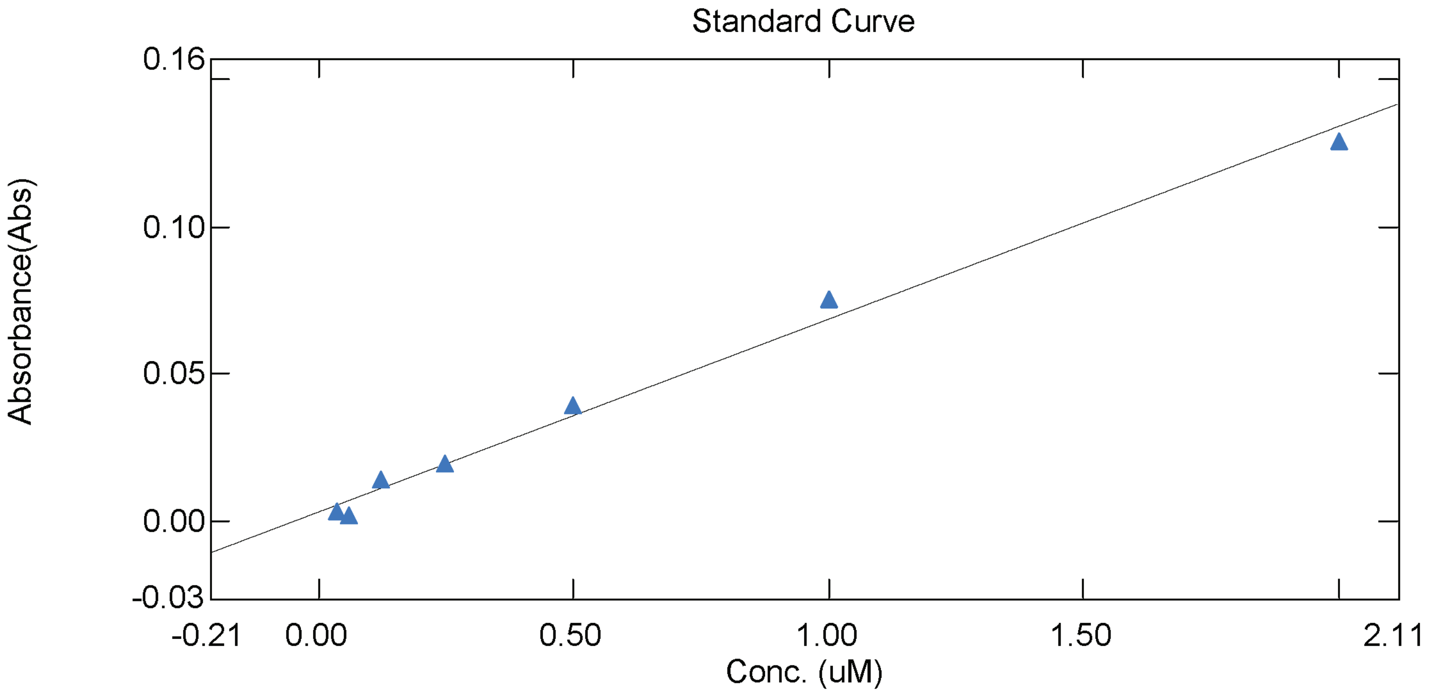


x = 15.2594 y - 0.0464410
r2 = 0.99068
r(m)2 = 0.99068

**a**

### Hydrodynamic Size in PBS
### NCL385-1 and NCL385-2

NCI Alliance for Nanotechnology in Cancer

**Low volume quartz cuvette, b = 10 mm, 25 °C, 633 nm laser λ, 173° scattering angle**
**Samples diluted 10 and 100-fold in PBS**

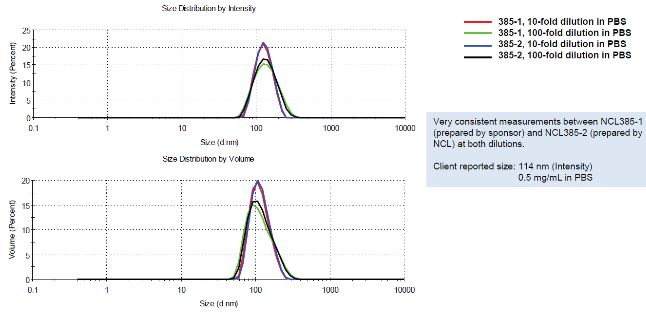


Very consistent measurements between NCL385-1 (prepared by sponsor) and NCL385-2 (prepared by NCL) at both dilutions.

Client reported size: 114 nm (Intensity)
0.5 mg/mL in PBS

| Sample | Sample concentration | Z-Avg, nm | PdI | Int-Peak, nm | %Int | Vol-Peak, nm | %Vol |
|---|---|---|---|---|---|---|---|
| NCL385-1 | 10-fold | 120.4 (1.1) | 0.041 (0.011) | 127.8 (1.0) | 100 (0) | 116.2 (1.4) | 100 (0) |
| " | 100-fold | 126.8 (1.8) | 0.127 (0.010) | 142.1 (3.2) | 100 (0) | 121.7 (2.8) | 100 (0) |
| NCL385-2 | 10-fold | 123.3 (0.8) | 0.042 (0.015) | 130.7 (1.0) | 100 (0) | 120.0 (1.0) | 100 (0) |
| " | 100-fold | 126.4 (2.6) | 0.094 (0. 29) | 139.6 (5.6) | 100 (0) | 122.2 (2.7) | 100 (0) |

**b**

### Hydrodynamic size in 10 mM NaCl
### NCL385-1

NCI Alliance for Nanotechnology in Cancer

**Low volume quartz cuvette, b = 10 mm, 25 °C, 633 nm laser l, 173° scattering angle**
**Samples diluted 10 and 100-fold in 10 mM NaCl**

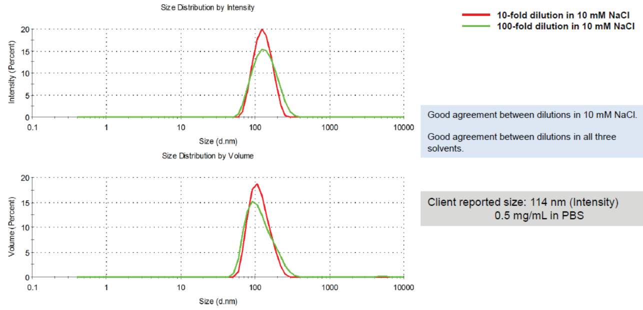


Good agreement between dilutions in 10 mM NaCl.

Good agreement between dilutions in all three solvents.

Client reported size: 114 nm (Intensity)
0.5 mg/mL in PBS

| Sample | Sample concentration | Z-Avg, nm | PdI | Int-Peak, nm | %Int | Vol-Peak, nm | %Vol |
|---|---|---|---|---|---|---|---|
| NCL385-1 | 10-fold | 121.4 (0.5) | 0.052 (0.013) | 130.0 (1.0) | 100 (0) | 117.2 (0.8) | 100 (0) |
| " | 100-fold | 126.8 (1.1) | 0.137 (0.015) | 140.9 (3.8) | 99.9 (0.5) | 120.7 (1.8) | 99.7 (1.0) |

Results are based on an average of 12 measurements.
Standard deviations are given in parentheses.

7

**c**

### Zeta Potential
### NCL385-2

NCI Alliance for Nanotechnology in Cancer

**Samples diluted 100-fold in 10 mM NaCl**
**Zeta potential measured at native pH/adjusted pH**
**25 °C, Applied Voltage: 151 V**

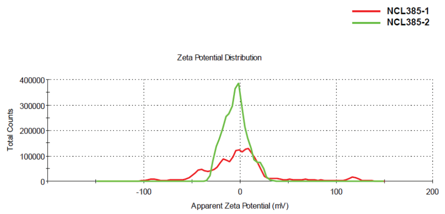


| Sample | pH | ZP (mV) |
|---|---|---|
| NCL385-1 | 7.24 | -1.1 (0.6) |
| NCL385-2 | 7.25 | -4.3 (1.4) |

Results are based on an average of 3 measurements

Sample shows neutral zeta potential in 10 mM NaCl, number of sub-runs increased for NCL385-2 analysis for better ZP measurement.

Client reported ZP: not reported

7

**d**

### Sterility & Endotoxin
### NCL385-1

NCI Alliance for Nanotechnology in Cancer

| Sample | Endotoxin: Turbidity, EU/mg nanogel (% spike recovery) | Endotoxin: Chromogenic, EU/mg nanogel (% spike recovery) | Sterility |
|---|---|---|---|
| NCL385-1 | 1.5 (113) | 2.6 (75) | Negative |
| EL | 8.79* | | |

* This endotoxin limit is applicable at HED 0.57 mg nanogel/kg/h

- No bacterial contamination was detected
- Endotoxin was below the EL

4

**e**

**Hydrodynamic size by Wyatt Dynapro Dynamic Light Scattering (DLS)**

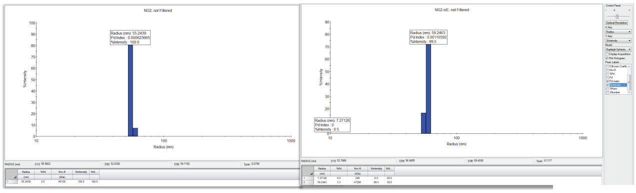

SANGs SANGs-siRNA EGFR

Representative DLS graphs showing hydrodynamic radius size measurement of SANGs and SANGs-siRNA EGFR in PBS, pH 7.4 @conc of 1 mg/ml SANGs

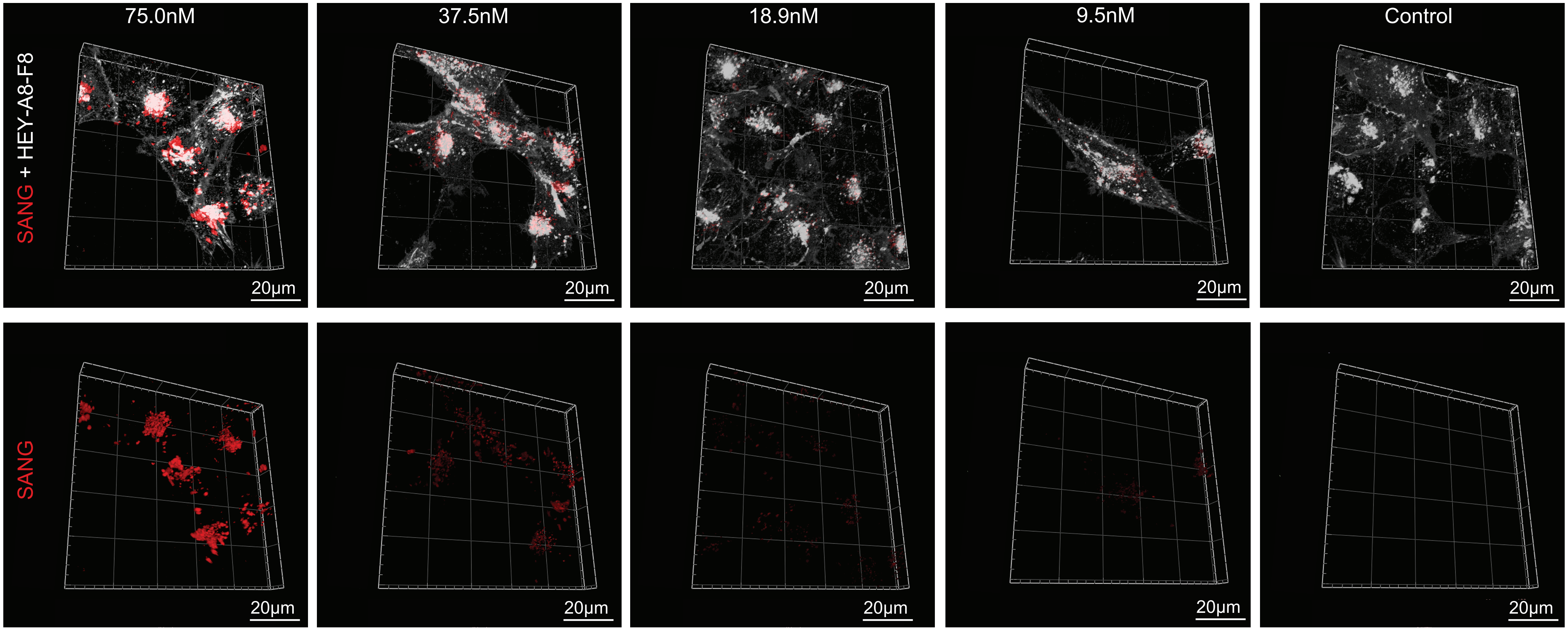

75.0nM
37.5nM
18.9nM
9.5nM
Control
SANG + HEY-A8-F8
SANG
20μm


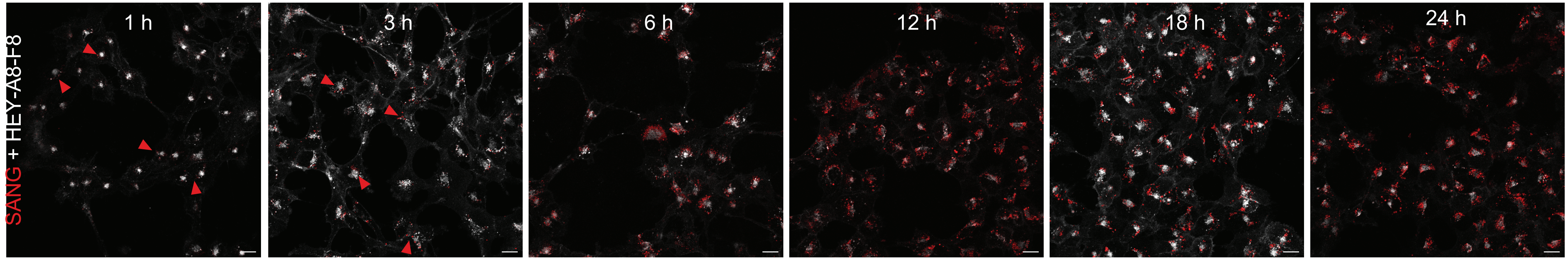

1 h
3 h
6 h
12 h
18 h
24 h
SANG + HEY-A8-F8

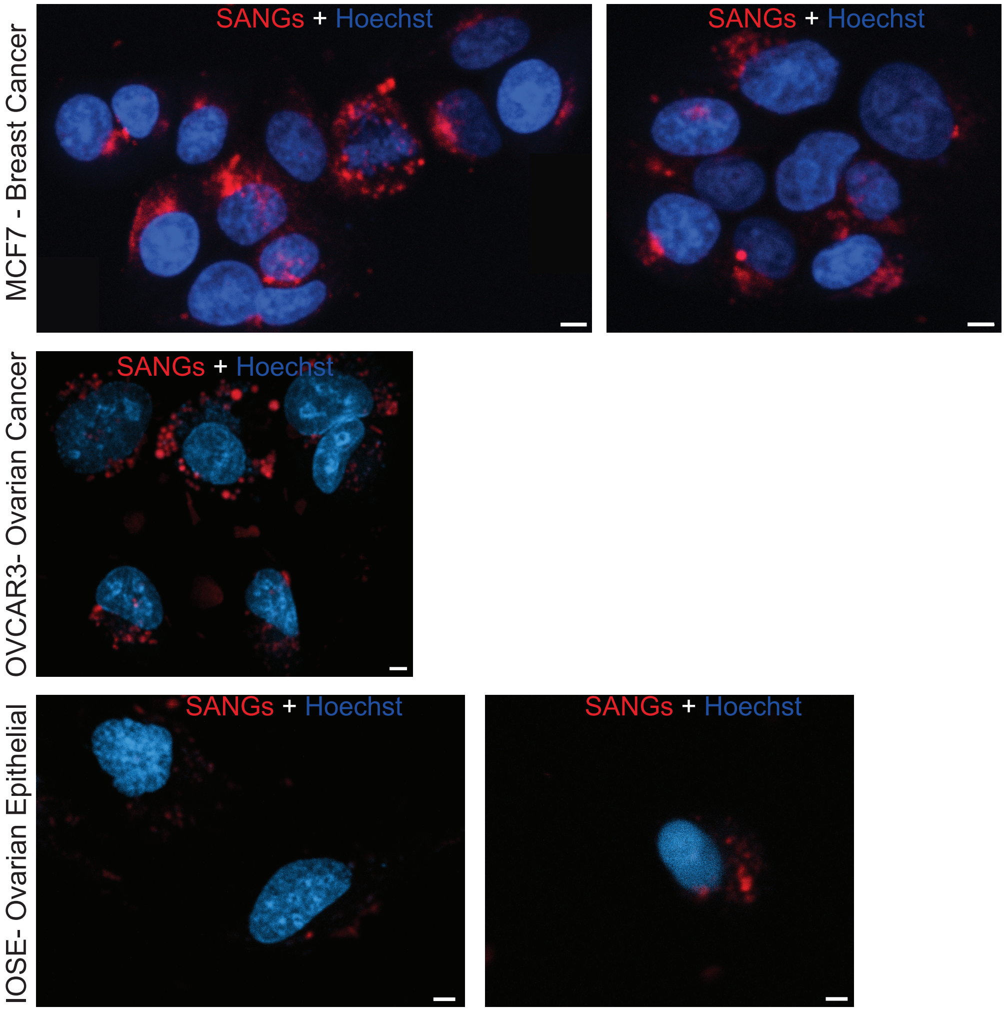
MCF7 - Breast Cancer
SANGs + Hoechst
SANGs + Hoechst
OVCAR3- Ovarian Cancer
SANGs + Hoechst
IOSE- Ovarian Epithelial
SANGs + Hoechst
SANGs + Hoechst

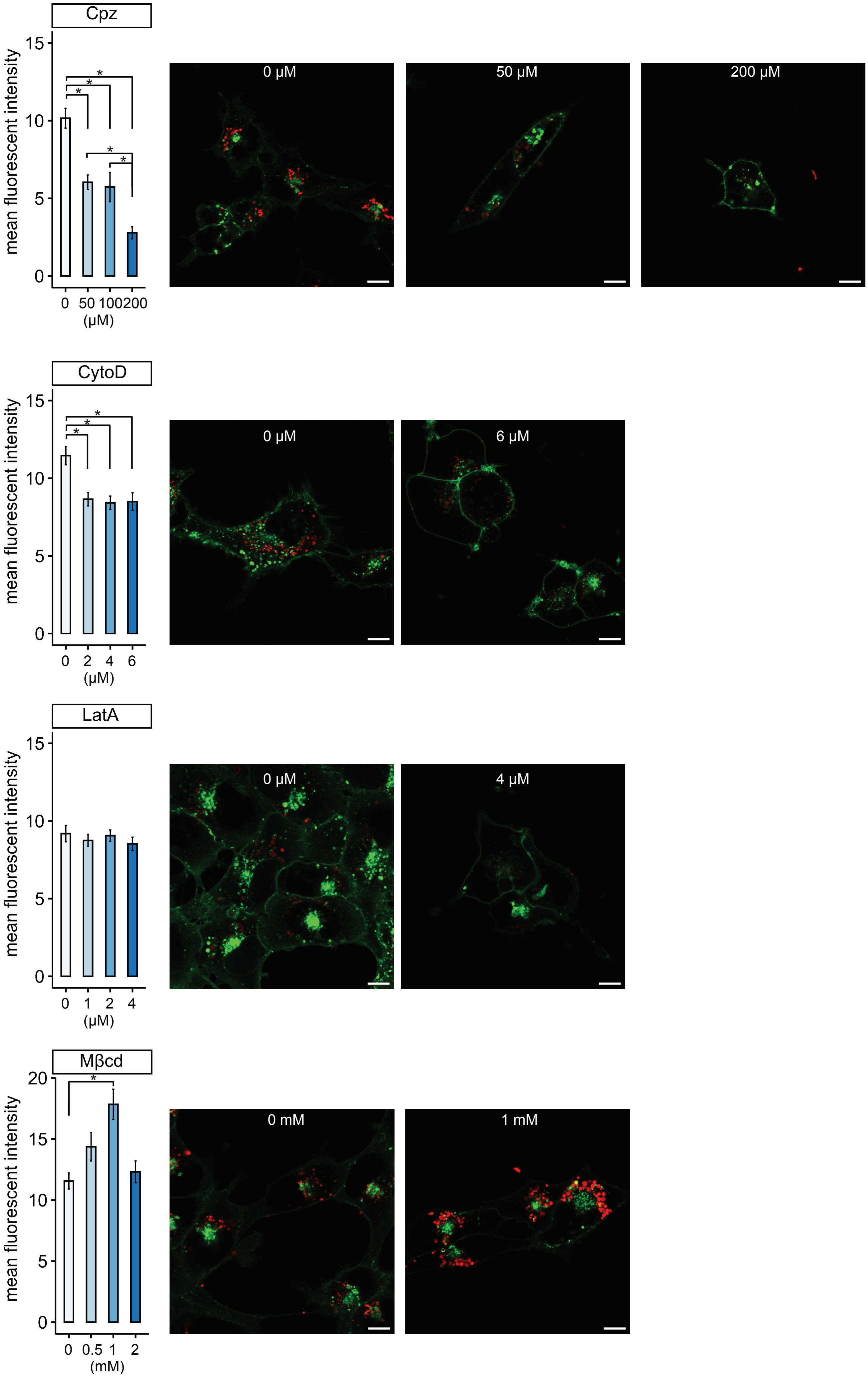
Cpz
mean fluorescent intensity
15
10
5
0
*
0
50
100
200
(μM)
0 μM
50 μM
200 μM
CytoD
mean fluorescent intensity
15
10
5
0
*
0
2
4
6
(μM)
0 μM
6 μM
LatA
mean fluorescent intensity
15
10
5
0
0
1
2
4
(μM)
0 μM
4 μM
Mβcd
mean fluorescent intensity
20
15
10
5
0
*
0
0.5
1
2
(mM)
0 mM
1 mM

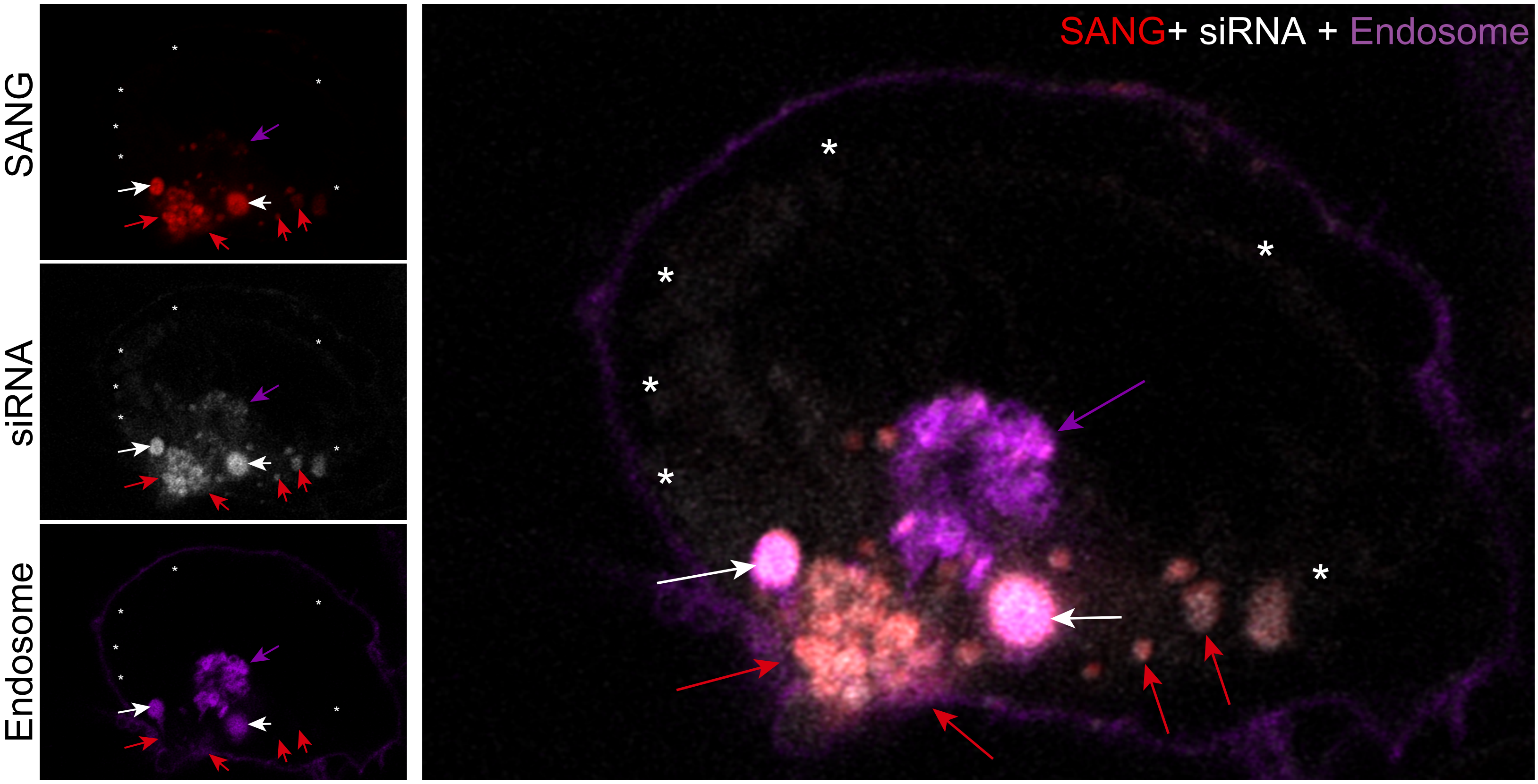
SANG
siRNA
Endosome
SANG+ siRNA + Endosome

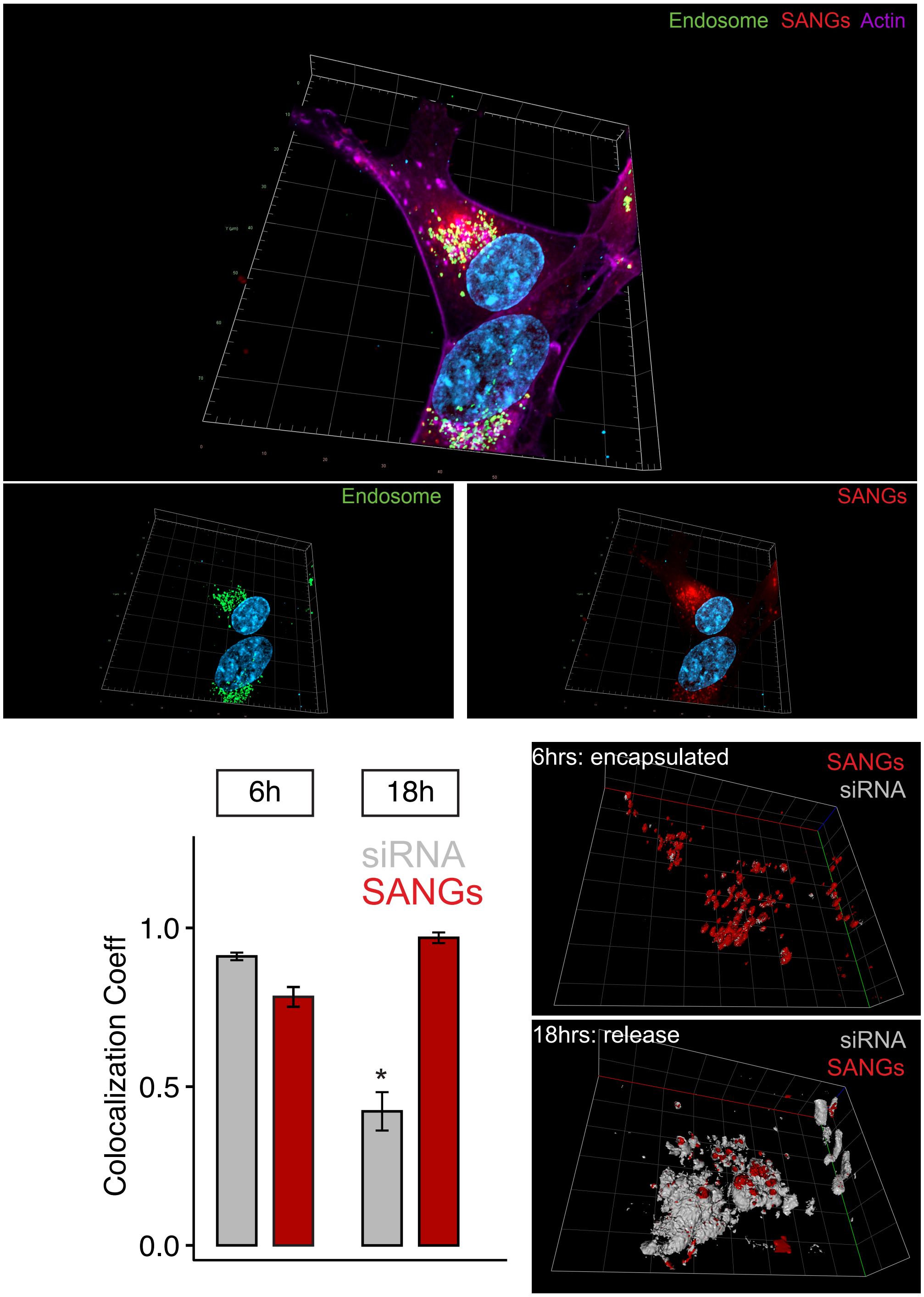
Endosome
SANGs
Actin
Endosome
SANGs
6h
18h
siRNA
SANGs
Colocalization Coeff
1.0
0.5
0.0
*
6hrs: encapsulated
SANGs
siRNA
18hrs: release
siRNA
SANGs

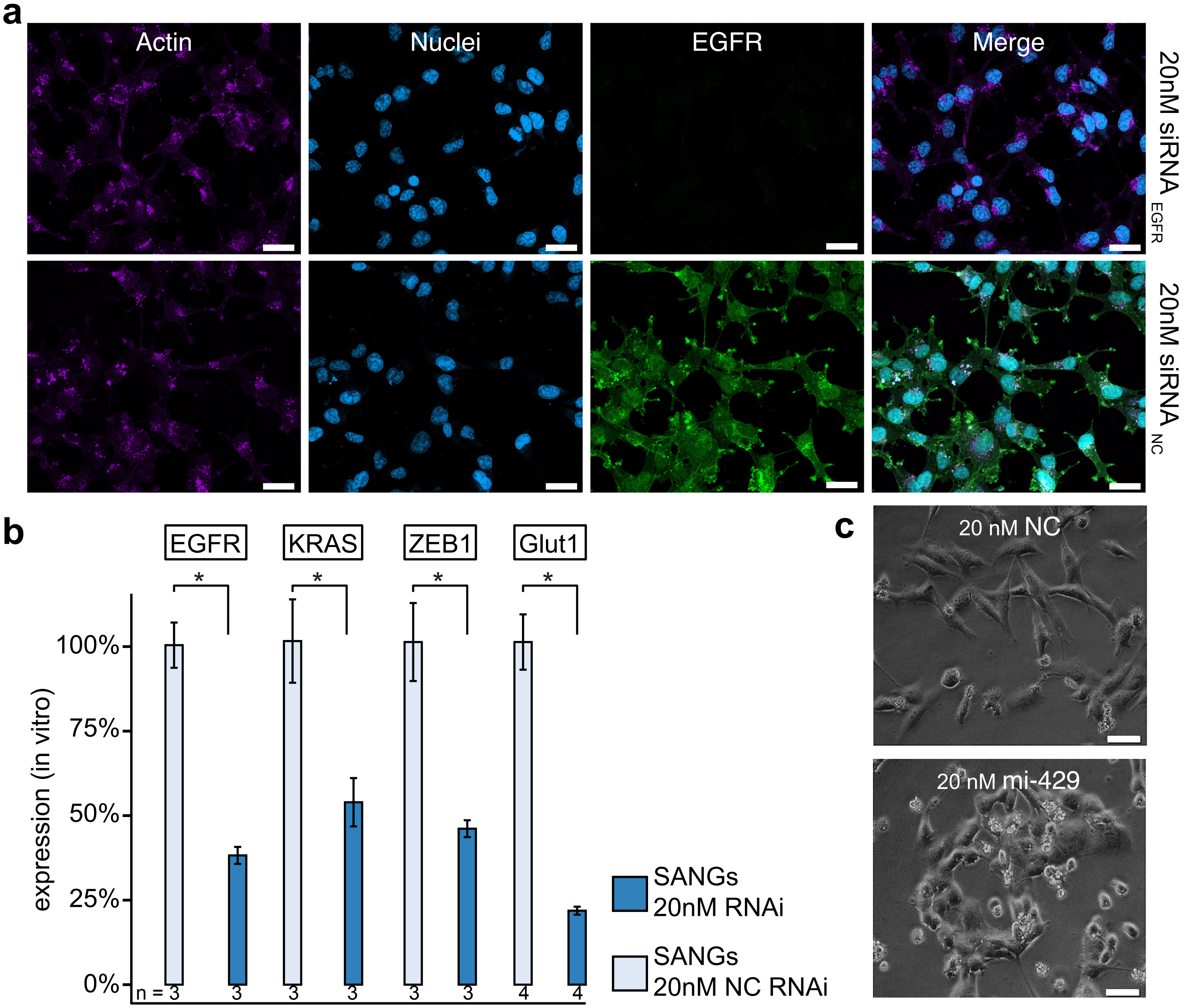
a
Actin
Nuclei
EGFR
Merge
20nM siRNA EGFR
20nM siRNA NC
b
EGFR
KRAS
ZEB1
Glut1
*
expression (in vitro)
100%
75%
50%
25%
0%
n = 3
3
3
3
3
3
4
4
SANGs
20nM RNAi
SANGs
20nM NC RNAi
c
20 nM NC
20 nM mi-429

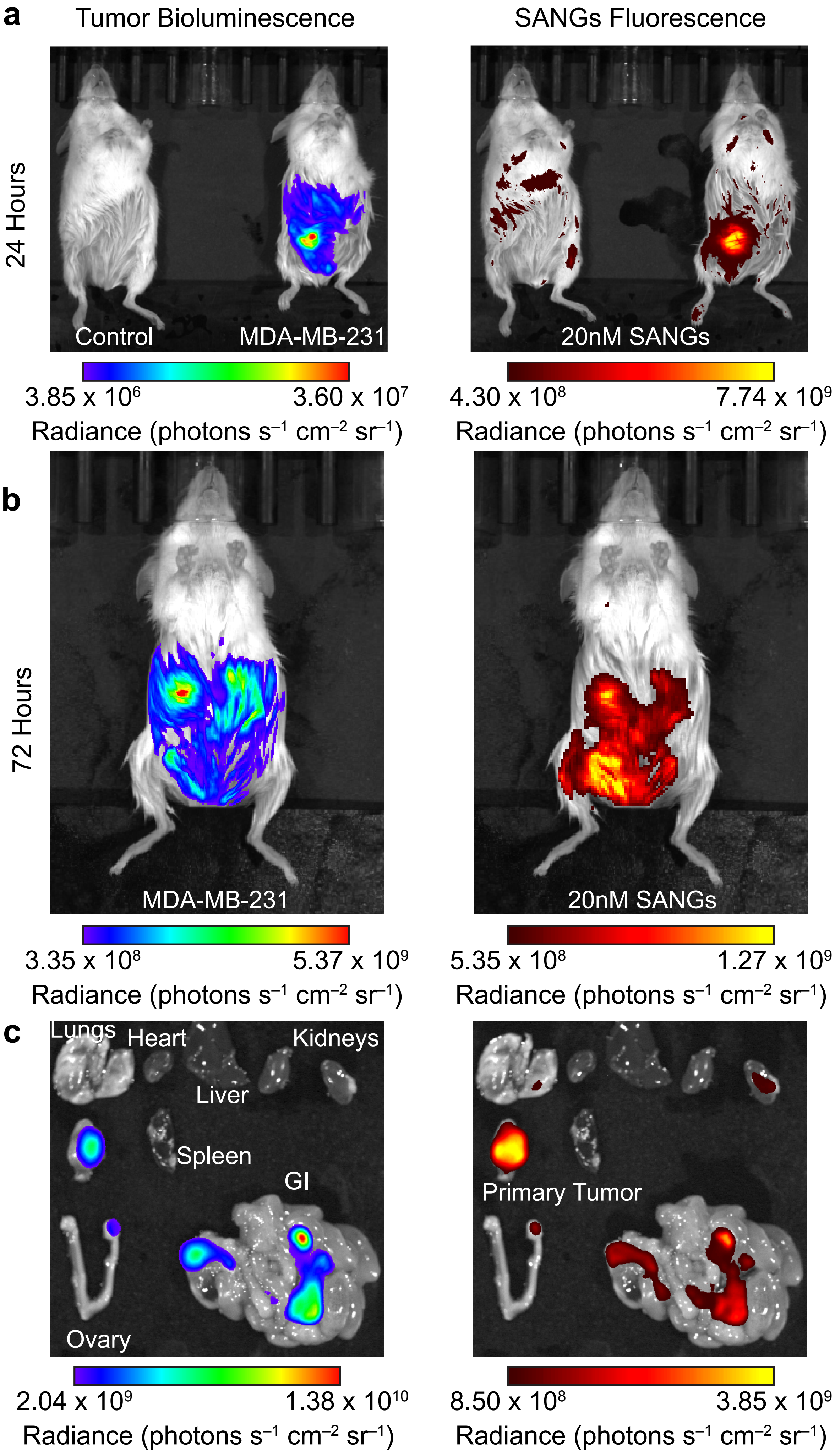
a
Tumor Bioluminescence
SANGs Fluorescence
24 Hours
Control
MDA-MB-231
20nM SANGs
3.85 x 10^6
3.60 x 10^7
Radiance (photons s^-1 cm^-2 sr^-1)
4.30 x 10^8
7.74 x 10^9
Radiance (photons s^-1 cm^-2 sr^-1)
b
72 Hours
MDA-MB-231
20nM SANGs
3.35 x 10^8
5.37 x 10^9
Radiance (photons s^-1 cm^-2 sr^-1)
5.35 x 10^8
1.27 x 10^9
Radiance (photons s^-1 cm^-2 sr^-1)
c
Lungs
Heart
Kidneys
Liver
Spleen
GI
Ovary
Primary Tumor
2.04 x 10^9
1.38 x 10^10
Radiance (photons s^-1 cm^-2 sr^-1)
8.50 x 10^8
3.85 x 10^9
Radiance (photons s^-1 cm^-2 sr^-1)

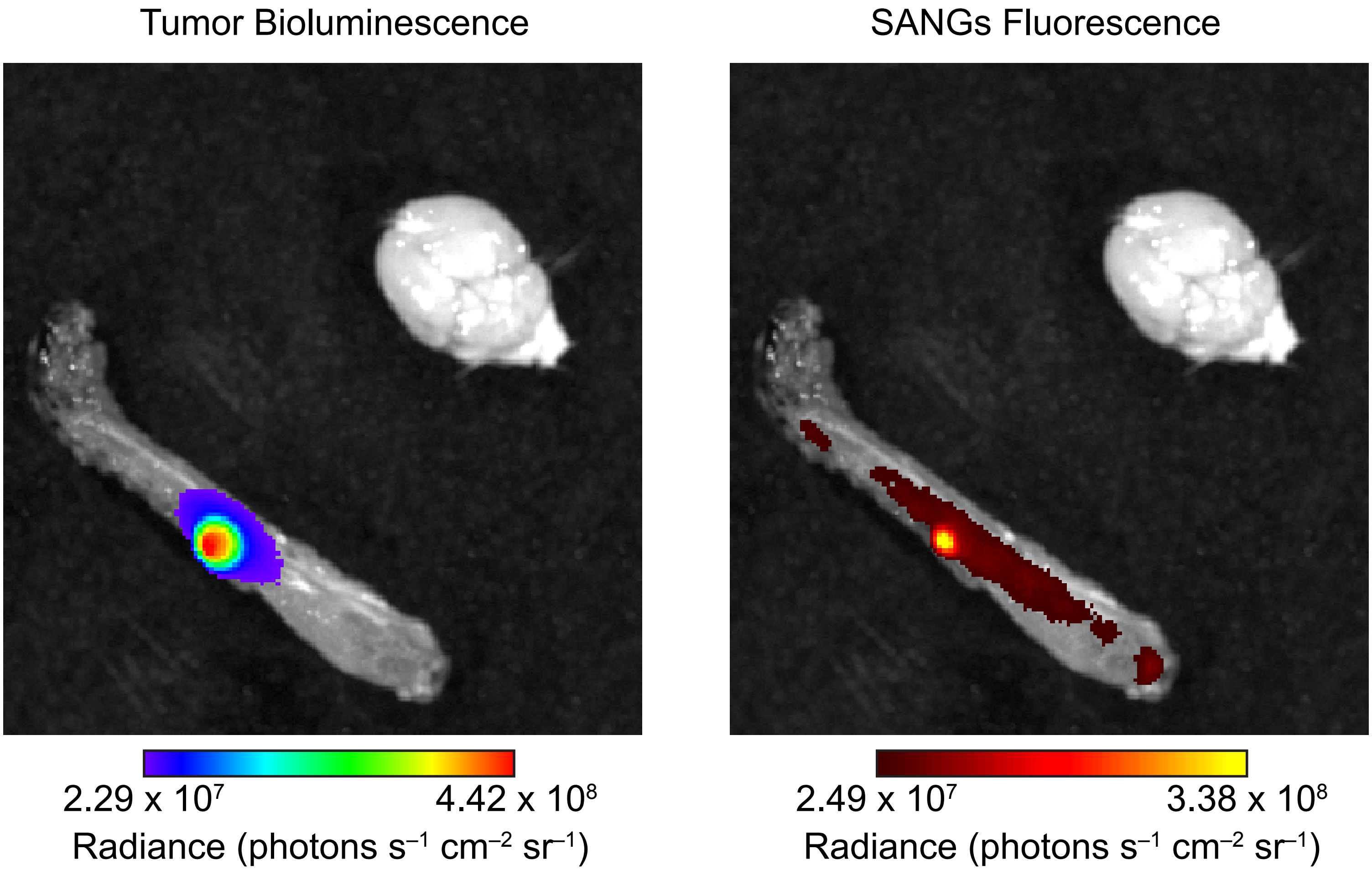
Tumor Bioluminescence
SANGs Fluorescence
2.29 x 10^7
4.42 x 10^8
Radiance (photons s^−1 cm^−2 sr^−1)
2.49 x 10^7
3.38 x 10^8
Radiance (photons s^−1 cm^−2 sr^−1)

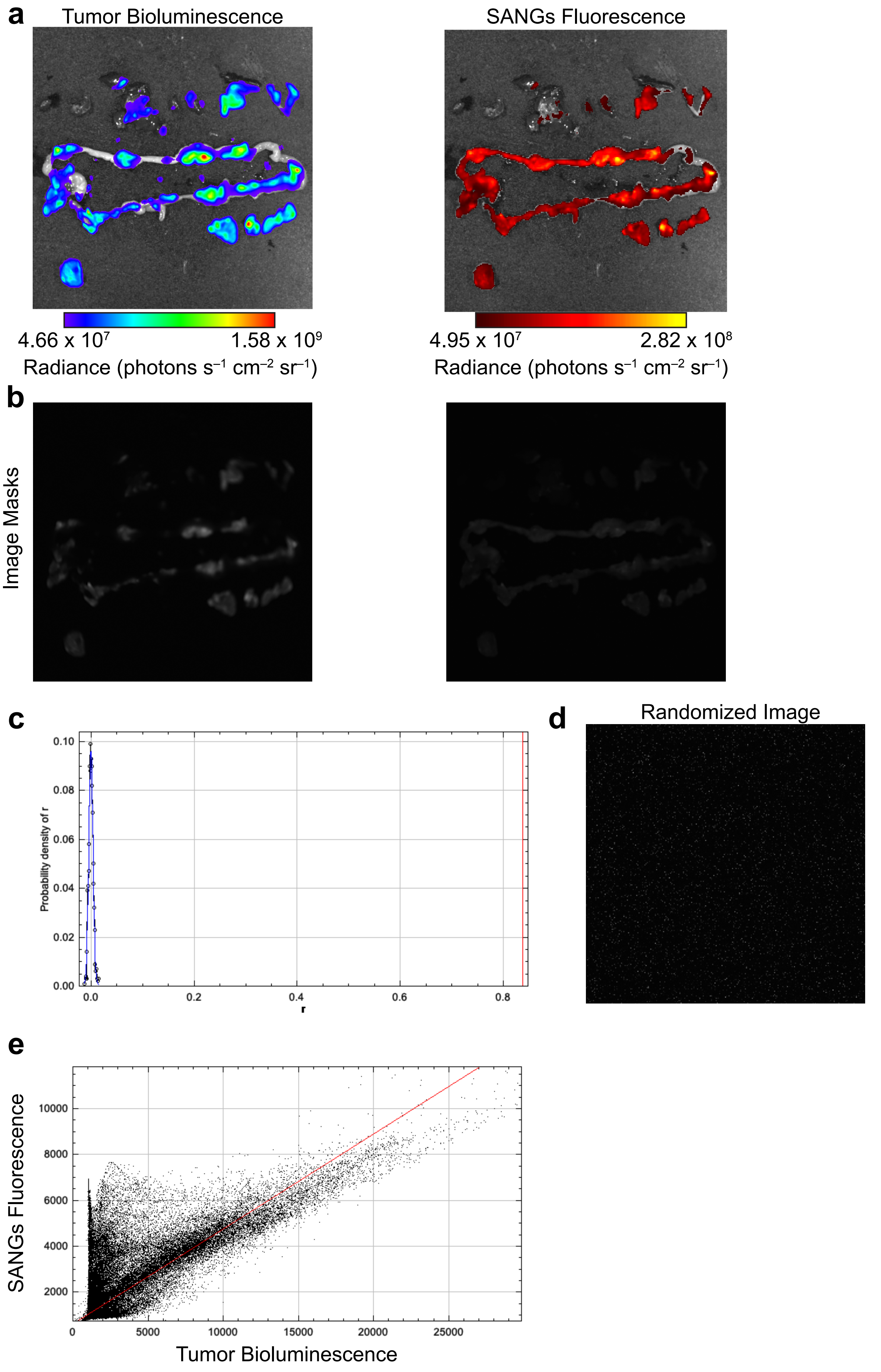

a
Tumor Bioluminescence
SANGs Fluorescence
4.66 x 10^7
1.58 x 10^9
Radiance (photons s^-1 cm^-2 sr^-1)
4.95 x 10^7
2.82 x 10^8
Radiance (photons s^-1 cm^-2 sr^-1)
b
Image Masks
c
Probability density of r
r
d
Randomized Image
e
SANGs Fluorescence
Tumor Bioluminescence

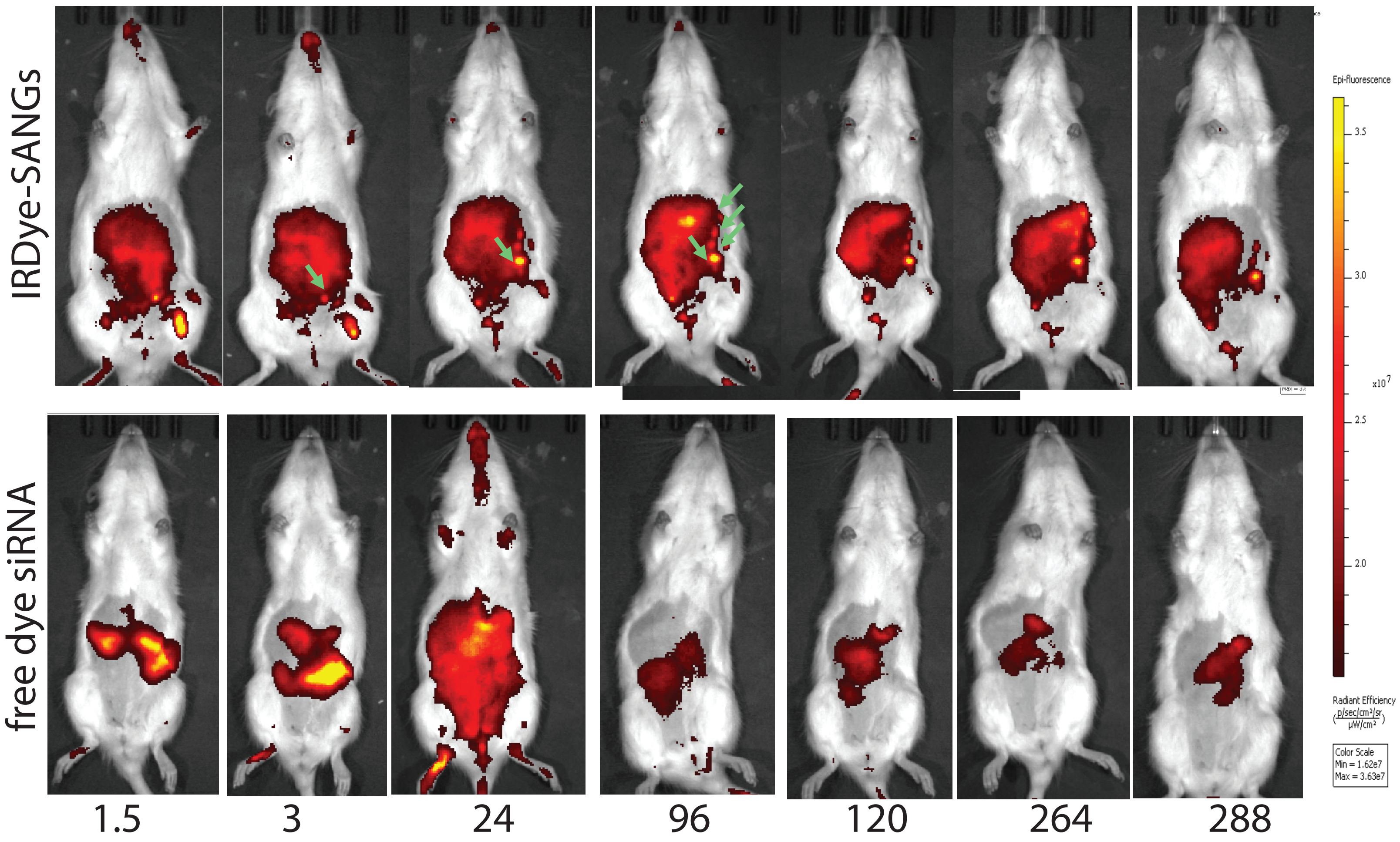
IRDye-SANGs
free dye siRNA
1.5
3
24
96
120
264
288
Epi-fluorescence
3.5
3.0
2.5
2.0
x10^7
Radiant Efficiency
(p/sec/cm²/sr / µW/cm²)
Color Scale
Min = 1.62e7
Max = 3.63e7

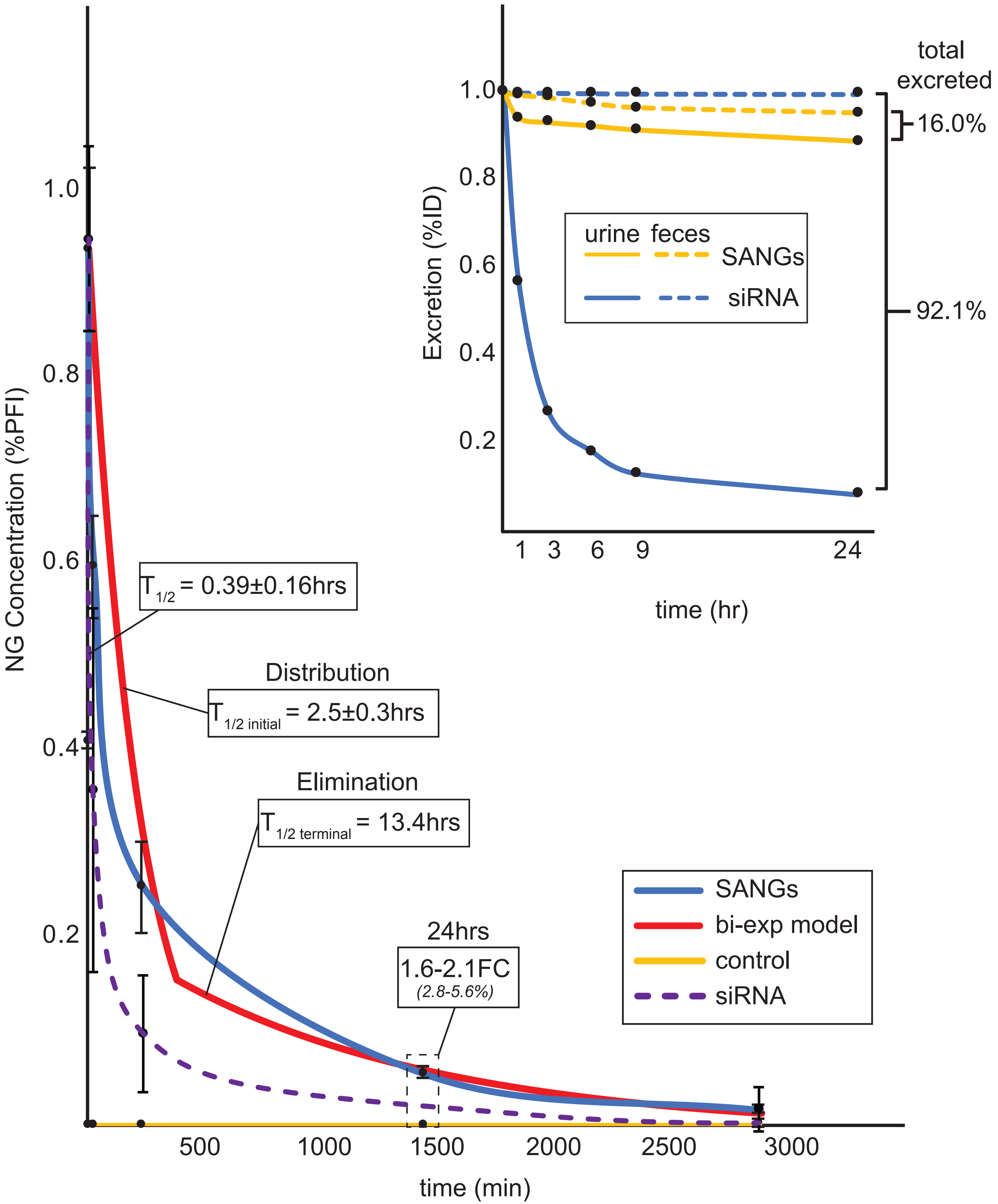

NG Concentration (%PFI)
1.0
0.8
0.6
0.4
0.2
500
1000
1500
2000
2500
3000
time (min)
$T_{1/2}$ = 0.39±0.16hrs
Distribution
$T_{1/2\ initial}$ = 2.5±0.3hrs
Elimination
$T_{1/2\ terminal}$ = 13.4hrs
24hrs
1.6-2.1FC
(2.8-5.6%)
SANGs
bi-exp model
control
siRNA
Excretion (%ID)
1.0
0.8
0.6
0.4
0.2
1
3
6
9
24
time (hr)
urine feces
SANGs
siRNA
total
excreted
16.0%
92.1%

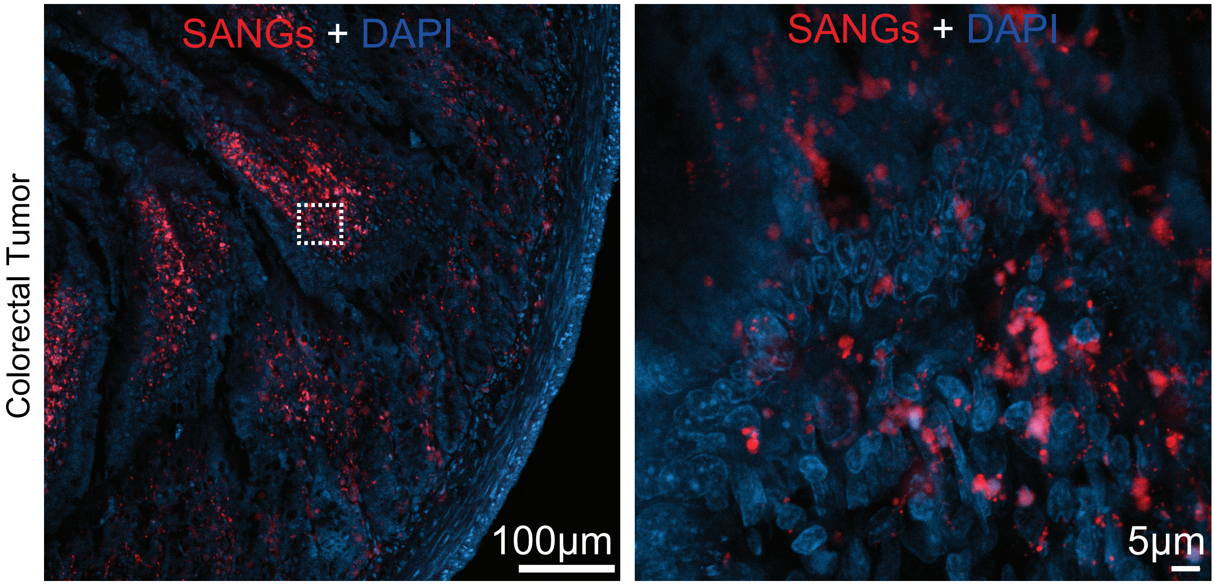
Colorectal Tumor
SANGs + DAPI
100μm
SANGs + DAPI
5μm

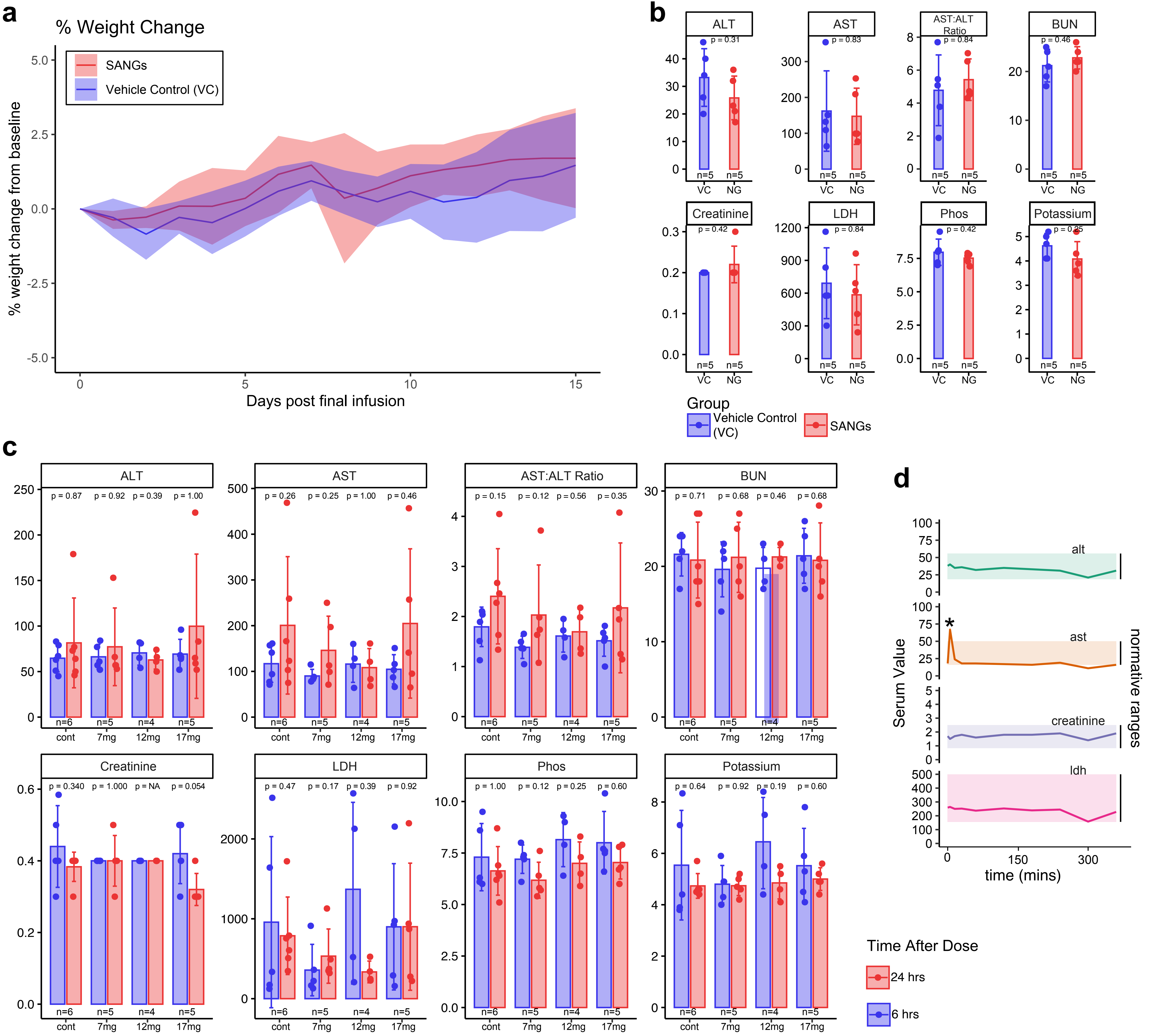
a
% Weight Change
SANGs
Vehicle Control (VC)
% weight change from baseline
Days post final infusion
b
ALT
AST
AST:ALT Ratio
BUN
Creatinine
LDH
Phos
Potassium
VC
NG
n=5
Group
Vehicle Control (VC)
SANGs
c
ALT
AST
AST:ALT Ratio
BUN
Creatinine
LDH
Phos
Potassium
cont
7mg
12mg
17mg
d
alt
ast
creatinine
ldh
Serum Value
normative ranges
time (mins)
Time After Dose
24 hrs
6 hrs

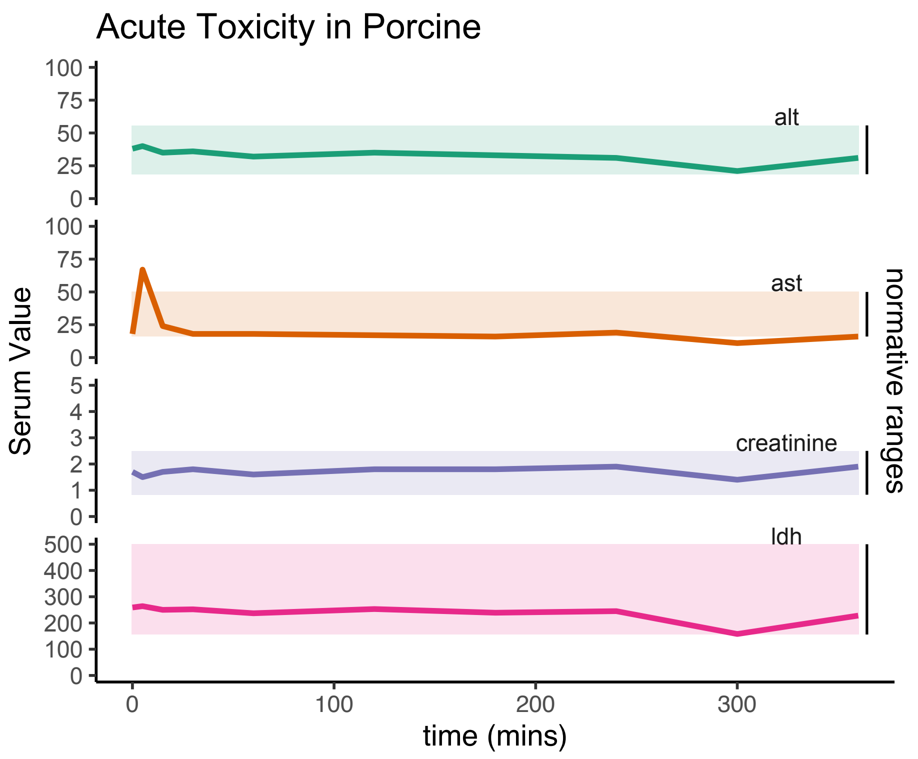
Acute Toxicity in Porcine
alt
ast
creatinine
ldh
Serum Value
normative ranges
time (mins)

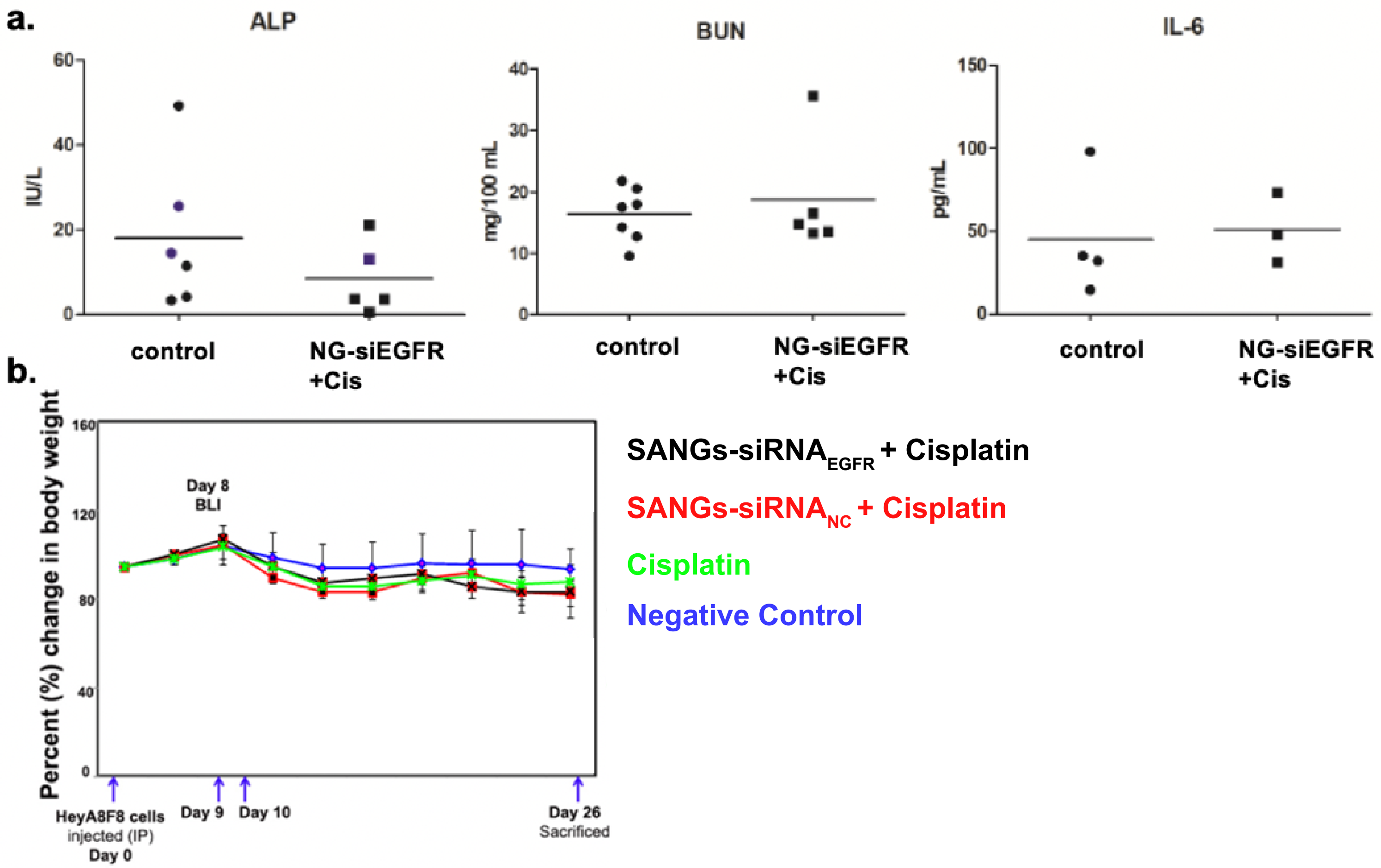

a.
ALP
IU/L
0
20
40
60
control
NG-siEGFR
+Cis
BUN
mg/100 mL
0
10
20
30
40
control
NG-siEGFR
+Cis
IL-6
pg/mL
0
50
100
150
control
NG-siEGFR
+Cis
b.
Percent (%) change in body weight
0
40
80
120
160
Day 8
BLI
HeyA8F8 cells
injected (IP)
Day 0
Day 9
Day 10
Day 26
Sacrificed
SANGs-siRNA$_{EGFR}$ + Cisplatin
SANGs-siRNA$_{NC}$ + Cisplatin
Cisplatin
Negative Control

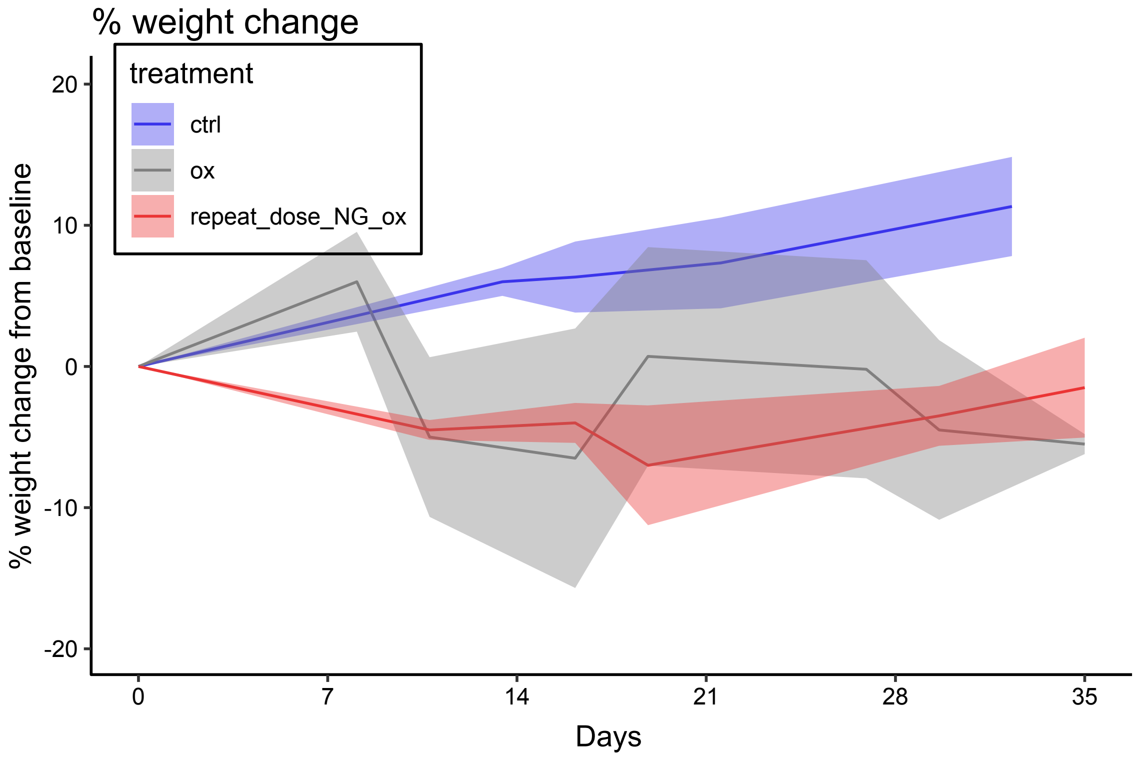

% weight change
treatment
ctrl
ox
repeat_dose_NG_ox
% weight change from baseline
20
10
0
-10
-20
0
7
14
21
28
35
Days

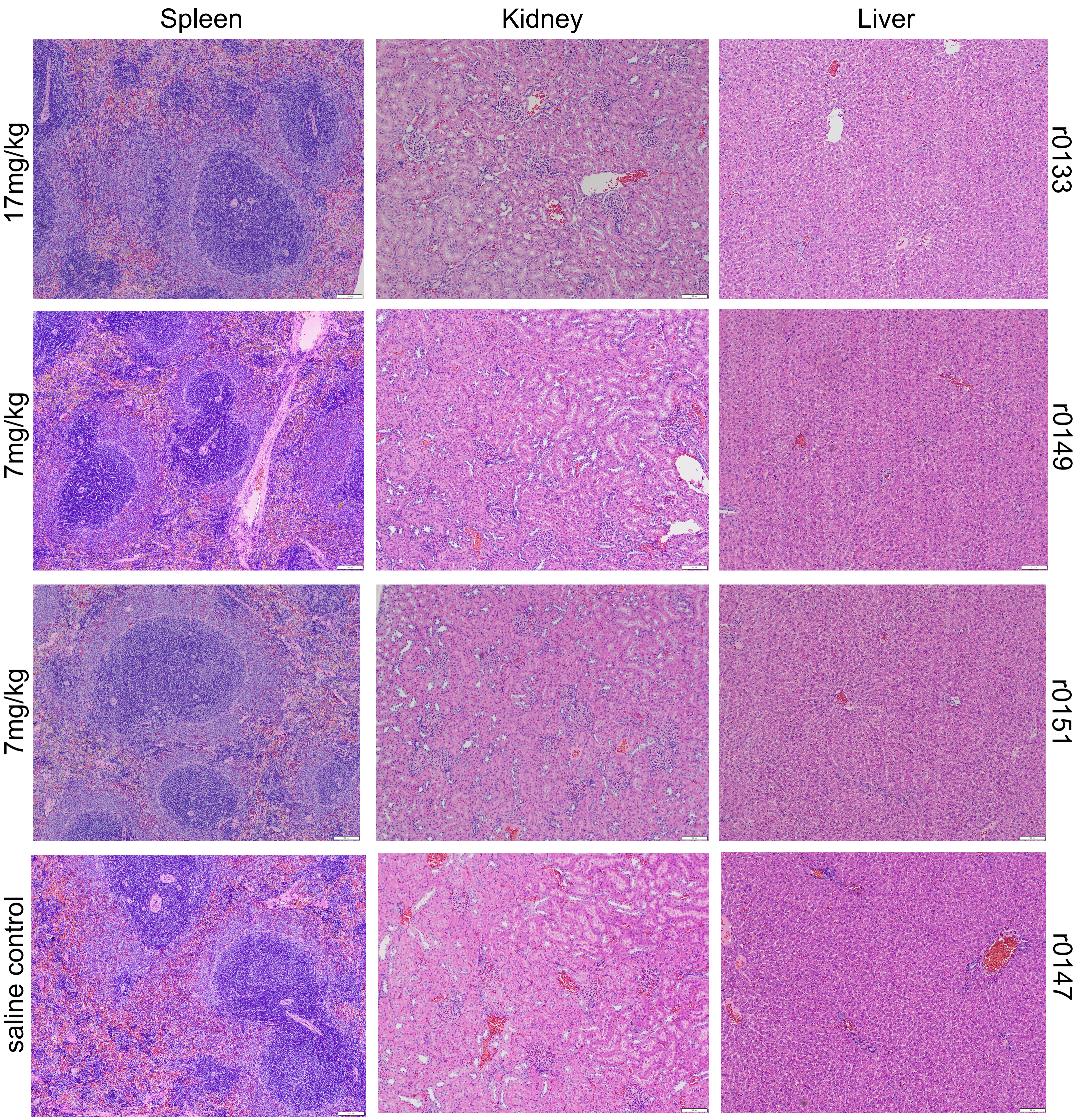

Spleen
Kidney
Liver
17mg/kg
r0133
7mg/kg
r0149
7mg/kg
r0151
saline control
r0147

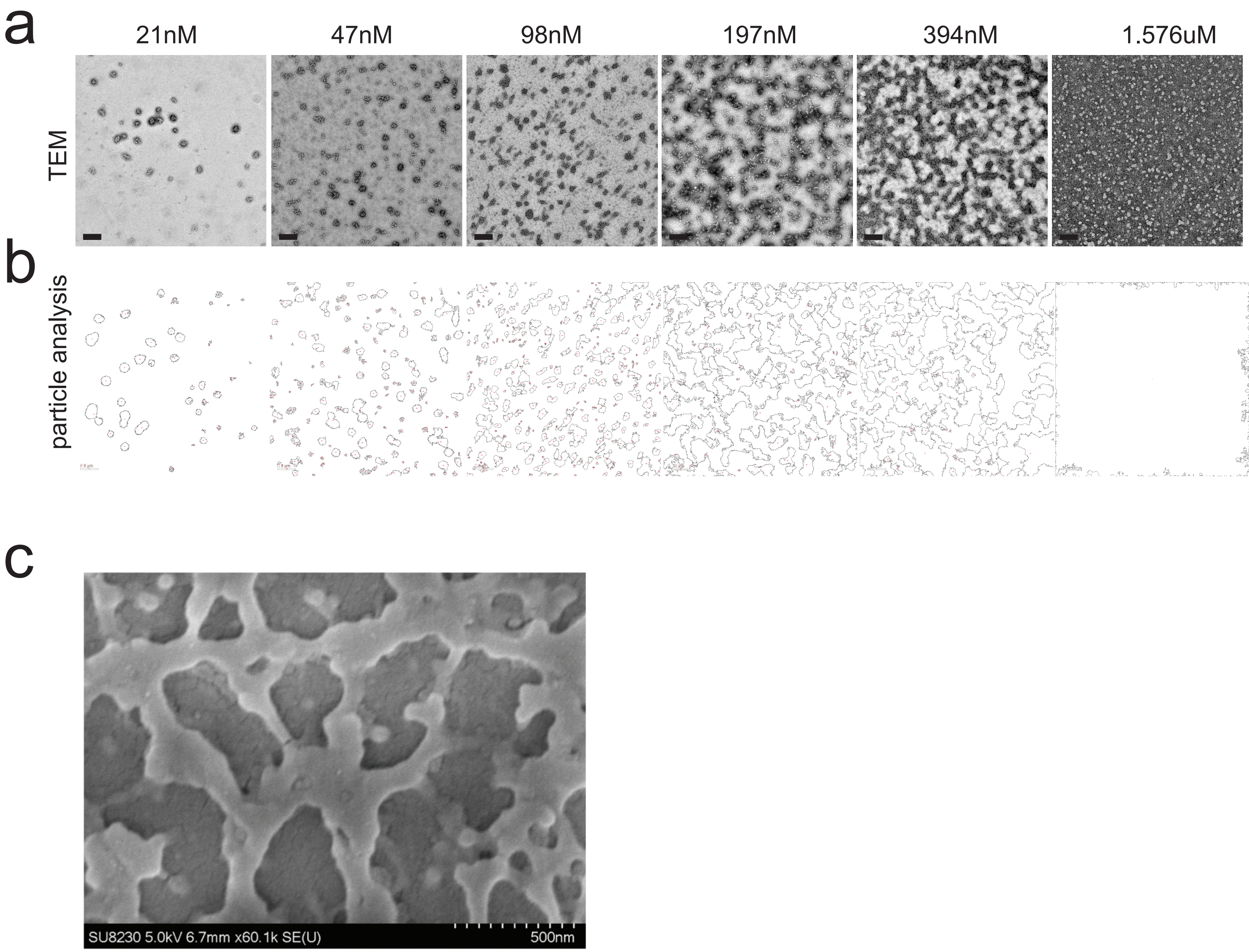
a
21nM
47nM
98nM
197nM
394nM
1.576uM
TEM
b
particle analysis
c
SU8230 5.0kV 6.7mm x60.1k SE(U)
500nm

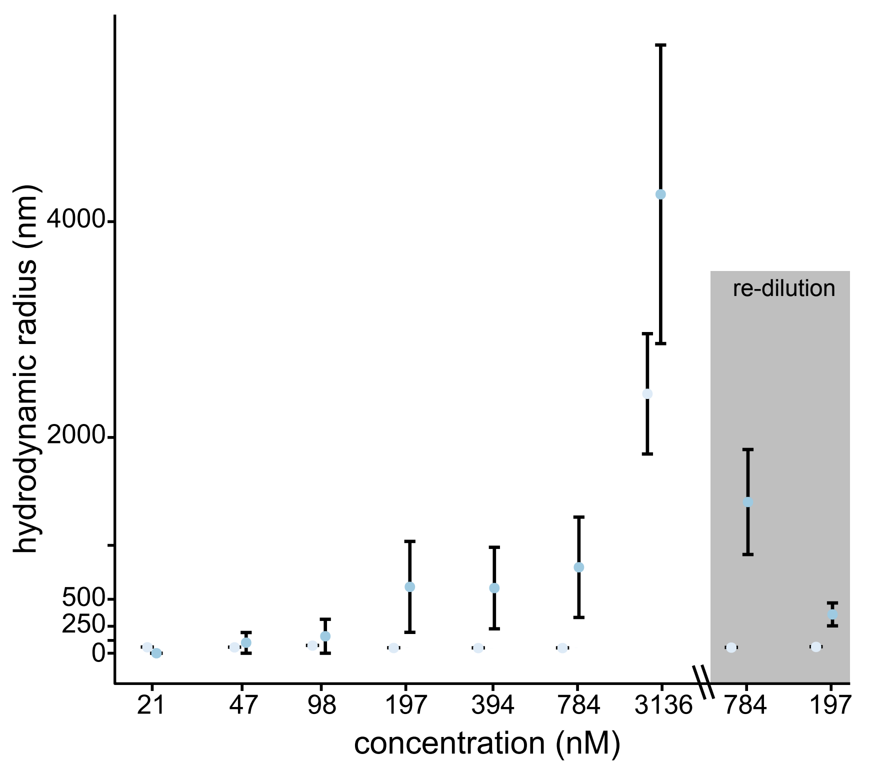

hydrodynamic radius (nm)
4000
2000
500
250
0
re-dilution
21
47
98
197
394
784
3136
784
197
concentration (nM)

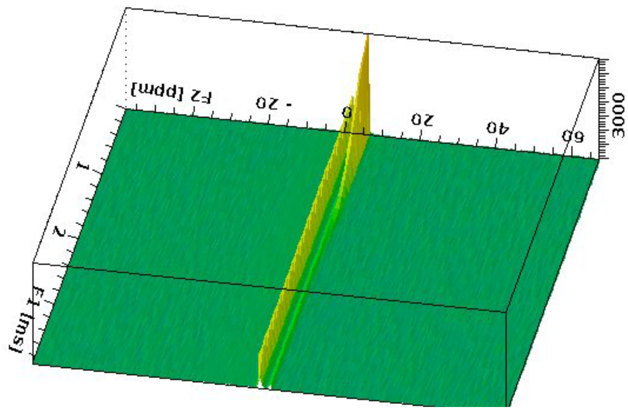

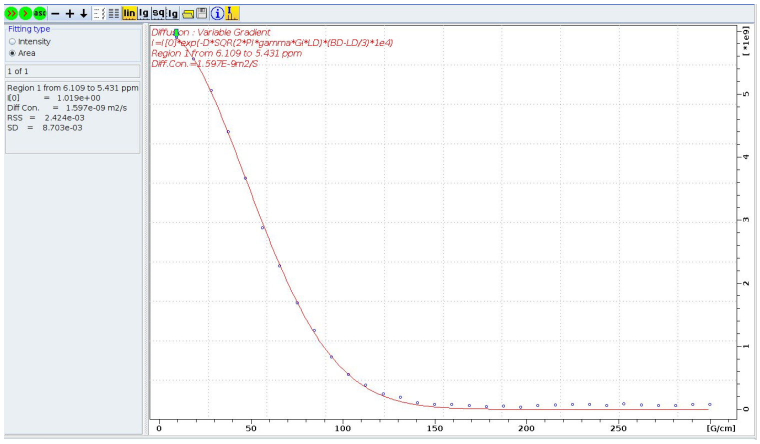
Fitting type
Intensity
Area
1 of 1
Region 1 from 6.109 to 5.431 ppm
I[0] = 1.019e+00
Diff Con. = 1.597e-09 m2/s
RSS = 2.424e-03
SD = 8.703e-03
Diffusion : Variable Gradient
I=I[0]*exp(-D*SQR(2*PI*gamma*Gi*LD)*(BD-LD/3)*1e4)
Region 1 from 6.109 to 5.431 ppm
Diff.Con. =1.597E-9m2/S
[G/cm]

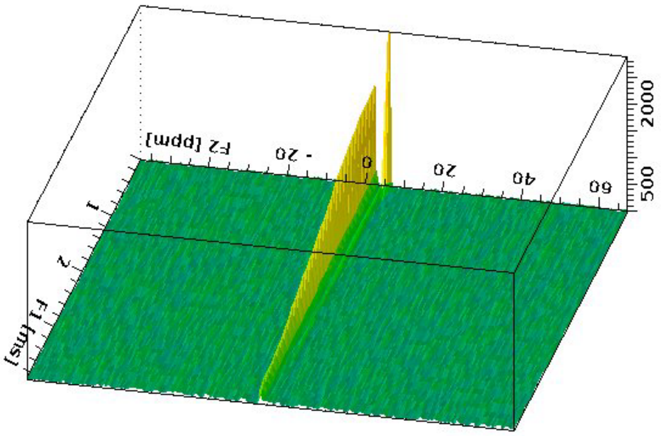

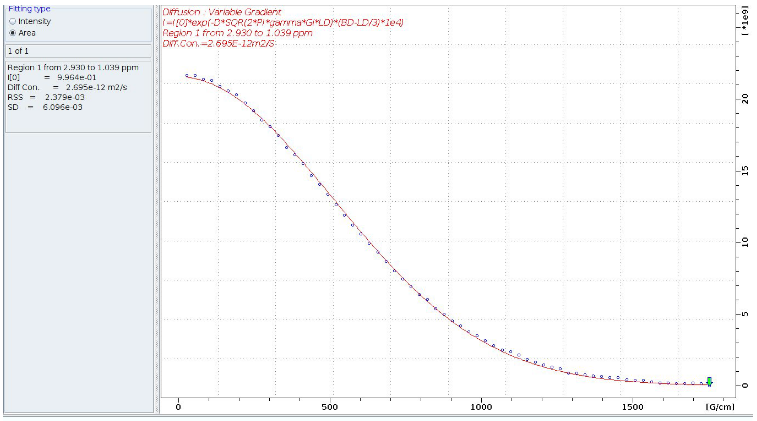
Fitting type
Intensity
Area
1 of 1
Region 1 from 2.930 to 1.039 ppm
I[0] = 9.964e-01
Diff Con. = 2.695e-12 m2/s
RSS = 2.379e-03
SD = 6.096e-03
Diffusion : Variable Gradient
I=I[0]*exp(-D*SQR(2*PI*gamma*Gi*LD)*(BD-LD/3)*1e4)
Region 1 from 2.930 to 1.039 ppm
Diff.Con. =2.695E-12m2/S
[G/cm]

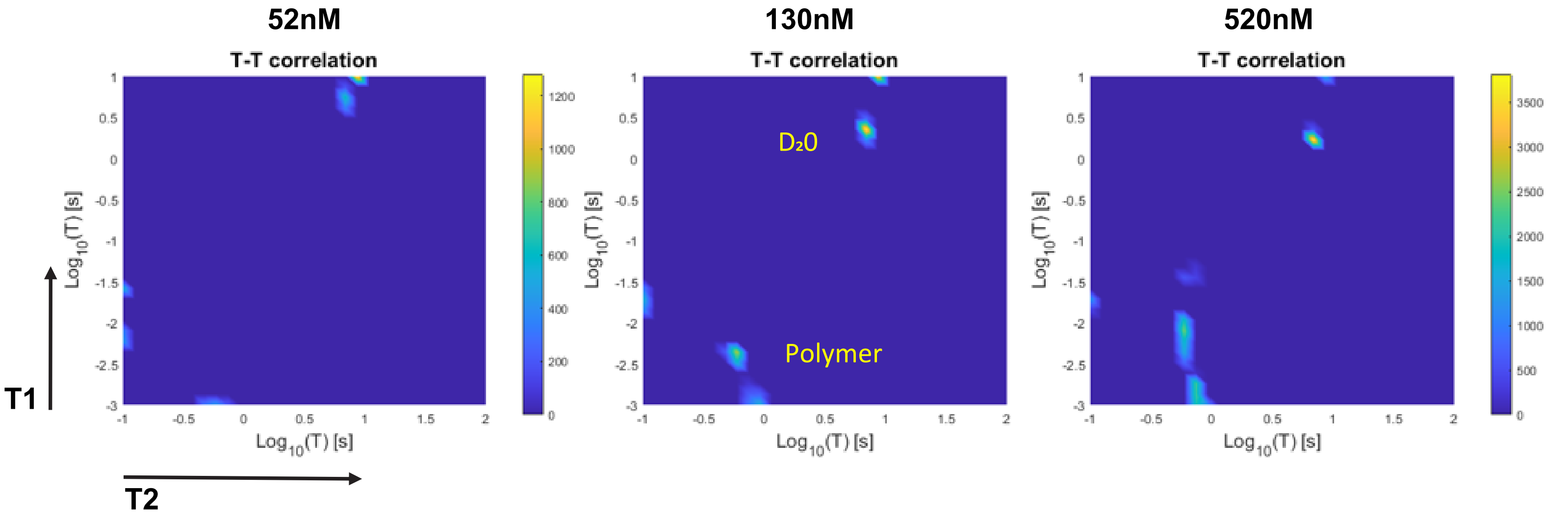
52nM
130nM
520nM
T-T correlation
T-T correlation
T-T correlation
D₂0
Polymer
Log10(T) [s]
T1
T2
